# Stochastic growth and selective stabilization generate stereotyped dendritic arbors

**DOI:** 10.1101/2024.05.08.591205

**Authors:** Rebecca Shi, Xue Yan Ho, Li Tao, Caitlin A. Taylor, Ting Zhao, Wei Zou, Malcolm Lizzappi, Kelsie Eichel, Kang Shen

## Abstract

Stereotyped dendritic arbors are shaped by dynamic and stochastic growth during neuronal development. It remains unclear how guidance receptors and ligands coordinate branch dynamic growth, retraction, and stabilization to specify dendritic arbors. We previously showed that extracellular ligand SAX-7/LICAM dictates the shape of the PVD sensory neuron via binding to the dendritic guidance receptor DMA-1, a single transmembrane adhesion molecule. Here, we perform structure-function analyses of DMA-1 and unexpectedly find that robust, stochastic dendritic growth does not require ligand-binding. Instead, ligand-binding inhibits growth, prevents retraction, and specifies arbor shape. Furthermore, we demonstrate that dendritic growth requires a pool of ligand-free DMA-1, which is maintained by receptor endocytosis and reinsertion to the plasma membrane via recycling endosomes. Mutants defective of DMA-1 endocytosis show severely truncated dendritic arbors. We present a model in which ligand-free guidance receptor mediates intrinsic, stochastic dendritic growth, while extracellular ligands instruct dendrite shape by inhibiting growth.

## Introduction

Neurons are the structural and functional units of the nervous system. Neurons form highly branched dendritic and axonal arbors that are crucial to establish neural circuits and appropriate receptive fields. The size of dendritic arbors is determined by cell-intrinsic mechanisms including transcription factors and cytoskeletal regulators^1,2^, cell-cell or cell-matrix interactions (guidance cues and receptors)^3^ and activity-dependent mechanisms^4,5^. Intracellularly, actin cytoskeleton is the major drive of dendrite and axonal morphogenesis^6^. Many actin regulators play critical roles in shaping dendritic branches, including the Rho family GTPases, Guanine Nucleotide exchange factors (GEFs), the WAVE regulatory complex (WRC), which coordinately control the activation of the ARP2,3 to promote the growth and branching of dendrites and dendritic spines^7,8^.

Extrinsic cues and their cell surface receptors guide dendritic morphogenesis. In the mammalian cortex, the growth direction of the apical dendrites of pyramidal neurons is guided by semaphorin 3A^9^. Classic axon guidance receptors Frazzled/DCC and ROBO mediate dendritic guidance at Drosophila midline^10^. The self-avoidance phenomenon, a critical contributor of dendritic arbor morphology, is mediated by homotypic cell adhesion receptors DSCAM and protocadherin^4,11–13^. How cell surface receptors activate signaling molecules to control cytoskeleton is not fully understood. It is generally assumed that ligand interaction with the extracellular domain of receptors transduces a signal to the cytoplasmic region to activate actin regulators.

Dendritic development, as revealed by imaging studies, is a dynamic and stochastic process. Individual developing dendrites cycle through dendritic initiation, growth and retraction to establish the shape of dendrites in many species^10,3,14^. Indeed, computational models can predict dendritic shape based on the dynamic features of branch tips and the rapid transitions between growth and retraction^15^. However, we do not understand how guidance receptors and their extracellular ligands mediate these complex and dynamic growth processes to build mature arbors with stereotyped arbors.

To probe this question, we studied the *C. elegans* multimodal sensory neuron PVD, which has a large dendrite arbor with repetitive menorah-like structures^16^. PVD dendrites innervate the skin of the animal, allowing the detection of mechanical, acoustic, proprioceptive and temperature stimuli^17^. The PVD dendrite starts to develop at the second larva stage (L2) and is fully formed in one day old adults (1DOA). During development, filopodia display continuous cycles of extension and retraction. Subsequent stabilization of the growing dendrites then establishes the mature dendrite morphology^14^.

The shape of PVD dendrites is determined by three extracellular cues (SAX-7, MNR-1, LECT-2) expressed by the target tissues and by a receptor complex (DMA-1 and HPO-30) on the dendrite. Among the ligands, SAX-7, the ortholog for the vertebrate L1CAM, is a transmembrane protein on skin cells and localizes to regularly spaced pattern^3,18,19^. The activation of DMA-1 requires all three ligands to specify the growth and branching of PVD dendrites in the extracellular space between the epidermis and muscle^3^. In PVD, DMA-1 binds to its coreceptor, a claudin homolog HPO-30, which is also required for PVD dendritic growth^20^; together this coreceptor complex recruits actin regulators, including TIAM-1 and the WAVE regulatory complex (WRC), to promote actin polymerization and dendritic growth^20–22^. Additionally, the furin-like proprotein convertase KPC-1 functions in PVD to promote dendritic growth^23–25^. KPC-1 was shown to downregulate overall protein levels of DMA-1^24,26^. However, it is unclear how KPC-1 regulates dendritic growth through its convertase activity and why increased levels of DMA-1 in *kpc-1(null)* leads to severe loss of PVD dendrites.

To understand how DMA-1 coordinates intracellular signaling and extracellular ligand interaction to translate stochastic dendritic growth into a stereotyped dendritic arbor shape, we performed genetic and cell biological experiments to dissect the structure and function of DMA-1. We reveal distinct roles for ligand-bound and ligand-free DMA-1. While ligand binding is required for DMA-1 to instruct the shape of dendritic arborization, it is not required for dynamic and stochastic dendritic growth. We observe that ligand binding to DMA-1 stabilizes the developing dendrites and instructs arbor shape. We also demonstrate that the extracellular ligand-binding domain and intracellular signaling domain of DMA-1 can function independently to stabilize the growing dendrite and to promote dynamic growth, respectively. Last, we show that DMA-1 is activated by KPC-1/Furin. KPC-1 cleaves the DMA-1 co-receptor HPO-30 and promotes DMA-1 internalization into endosomes and subsequent DMA-1 recycling to the plasma membrane. Together, the dual functions of this guidance receptor explain the dynamic, stochastic growth of the developing dendrites and the ultimate stereotypical arbor shape.

## Results

### Ligand-free DMA-1 receptor supports dendritic growth

PVD neurons have large and organized dendritic arbors, consisting of primary (1°), secondary (2°), tertiary (3°), and quaternary (4°) dendritic branches, resulting in numerous menorah-like structures (Figure 1A). We have previously shown that DMA-1 functions cell-autonomously to promote PVD dendritic formation and growth^1^, while its ligand SAX-7/L1CAM is required in neighboring epidermal cells to dictate the shape of the dendrites^3,18^. *dma-1(null)* mutants show dramatically reduced total dendritic length and completely lack menorah-like structures (Figure 1B and 1C). Interestingly, *sax-7* mutants does not phenocopy *dma-1(null)* mutants. While *sax-7(null)* mutants also lack menorahs, the total dendritic length is significantly more extensive than *dma-1(null)* mutants. We also observed that the dendritic arbors in *sax-7(null)* mutants are disorganized (Figure 1B and 1C). The disorganized arbors observed in *sax-7(null)* animals requires DMA-1 as we have previously shown that *dma-1(null); sax-7(null)* double mutants phenocopy *dma-1(null)* mutants, with few dendritic branches^3^. This indicates that DMA-1 continues to promote disorganized dendrite growth in the absence of SAX-7. We predict that DMA-1 plays distinct roles to promote growth (independent of SAX-7) and stabilization (dependent on SAX-7). Therefore, we hypothesize that distinct functional domains of DMA-1 – non-SAX-7 binding and SAX-7 binding domains, might be responsible to either promote growth or stabilization of dendrites.

**Figure 1.**
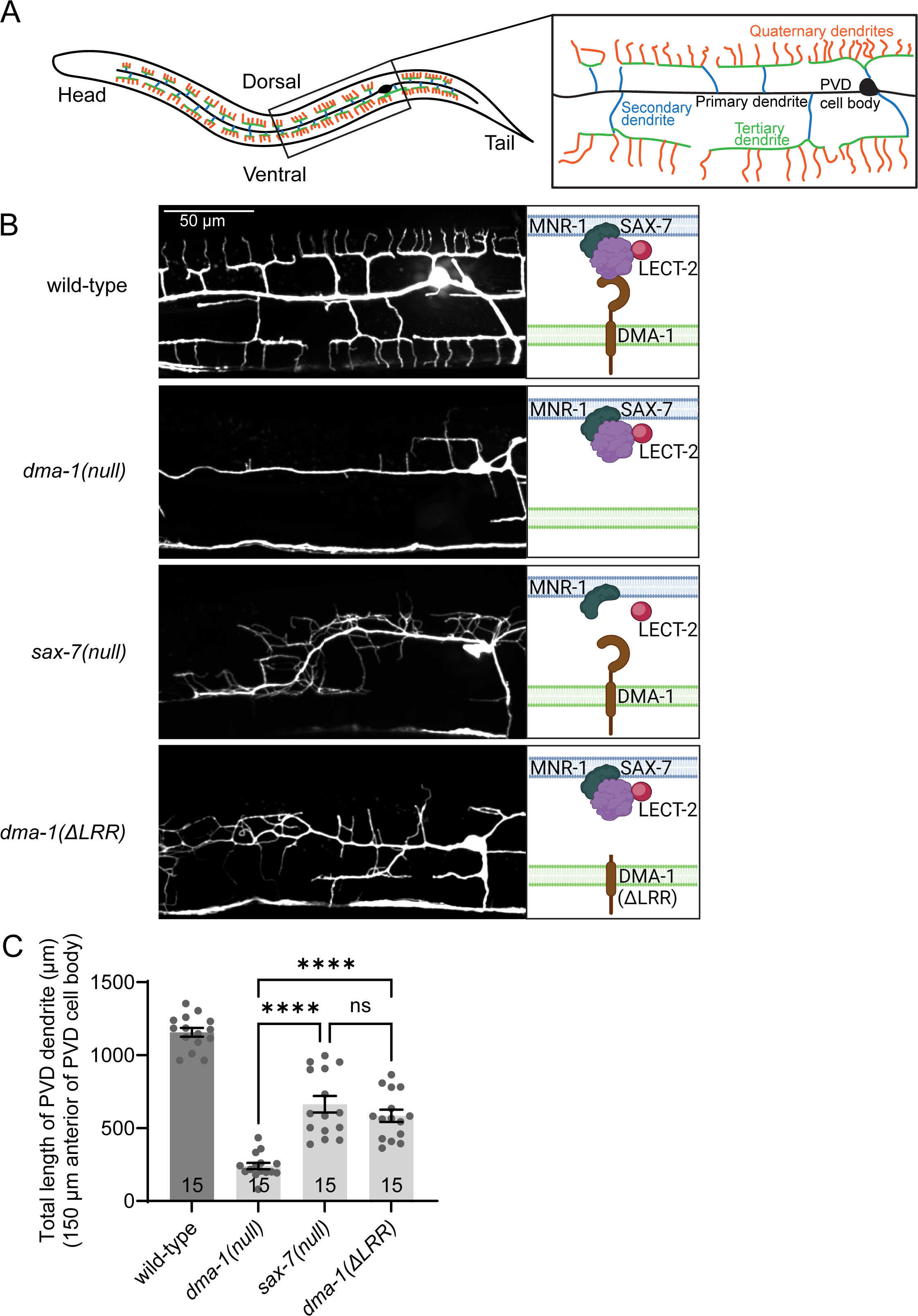
Ligand-free DMA-1 receptor supports dendritic growth. (A) Schematic of a WT animal with PVD neuron (left) and close-up of the boxed region (right). Anterior is left and ventral is down in this and all following images that shows PVD morphology. (B) Representative images of PVD dendrite morphology in wild-type, *dma-1(null), sax-7(null)* and *dma-1(ΔLRR)* mutant animals at 1DOA stage (left). Scale bar, 50 µm. Image representative of 15 animals. Schematic images of molecules that are present in the corresponding animals (right). The cell membrane is drawn in pink, and leaflets of the plasma membrane are differentiated in dark and light colors. MNR-1 is drawn in dark green. SAX-7 is drawn in purple. LECT-2 is drawn in red. DMA-1 is drawn in brown. (C) Quantification of the total length of PVD dendrite, 150 µm anterior to the PVD cell body. n values within each bar. Error bars indicate the standard error of proportion. Statistical comparison was performed using Brown-Forsythe ANOVA with Dunnett’s multiple comparisons test.

DMA-1 is a single transmembrane protein with an extracellular domain, a transmembrane and short intracellular region. The entire extracellular domain is predicted to contain a continuous leucine-rich-repeat domain (436 amino acid) and a 72 amino-acid juxtamembrane sequence. To differentiate between the extracellular and intracellular functions of DMA-1, we used CRISPR-Cas9 to precisely delete the entire leucine-rich-repeat domain of DMA-1 from its endogenous locus (*dma-1(*Δ*LRR)*). This truncated DMA-1 molecule retains the juxtamembrane sequence, as well as intact transmembrane and intracellular domains. As DMA-1 is a cell surface receptor that bridges extracellular cues with intracellular signaling, one would predict that removing the extracellular LRR domains (DMA-1(ΔLRR)) and hence its ability to bind its ligand, would abolish the function of DMA-1 and phenocopy *dma-1(null)* mutants. Interestingly, *dma-1(*Δ*LRR)* mutants displayed exuberant but disorganized arbors, indistinguishable from the *sax-7(null)* mutants (Figure 1B and 1C). Together these results suggest that even though the extracellular, ligand-binding domain of DMA-1 is required for establishing the arbor shape, the non-ligand-binding transmembrane and cytosolic domains of DMA-1 are likely to be sufficient to promote the growth and branching of dendrites.

### Ligand-bound DMA-1 receptor stabilizes dendritic branches

Dynamic imaging experiments from previous studies showed that developing dendrites are highly dynamic and undergo growth and retraction, while fully established dendrites are stabilized and much less dynamic^10^. PVD dendrites in wild-type animals are fully formed at the end of the L4 stage, and dendrites exhibit little continuous growth and retraction at this stage, thus exhibiting the typical transition from developmental dynamicity to stability (Figure 2A, 2B and 2C). We hypothesize that the transition from dynamic to stable dendrites might be impaired in *sax-7(null)* and *dma-1(*Δ*LRR)* animals, resulting in disorganized and extensive dendritic growth. To understand whether the disorganized dendritic arbors in *sax-7(null)* and *dma-1(*Δ*LRR)* animals are caused by defects in this transition, we quantified the growth and retraction of dendrites in wild-type, *sax-7(null)*, *dma-1(*Δ*LRR)* and *lect-2(null)* animals with time-lapse imaging. LECT-2 is one of the ligands, required for PVD dendritic shape^14,27,28^. Indeed, in *sax-7(null)*, *dma-1(*Δ*LRR)* and *lect-2(null)* mutants, dendrites exhibit continuous, dynamic growth and retraction (Figure 2A, 2B and 2C). These results indicate that ligand binding is required for the stabilization of the dendrites and could inhibit continuous dynamicity. This also further supports the notion that the intracellular domain of DMA-1 is sufficient to promote growth promoting activity.

**Figure 2.**
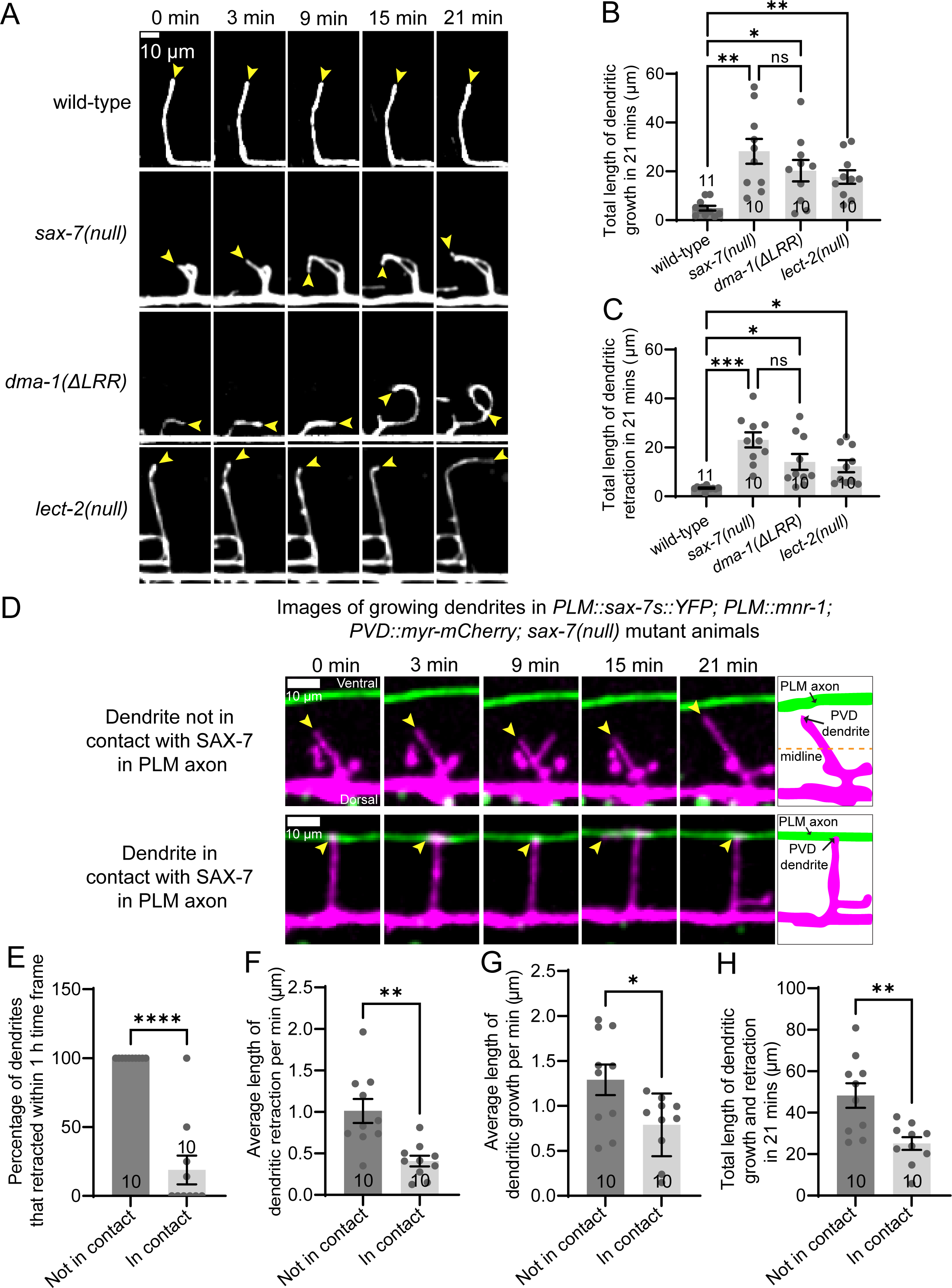
Ligand-bound DMA-1 receptor inhibits dendritic growth. (A) Time-lapse imaging of developing PVD secondary dendrites in wild-type, *sax-7(null), dma-1(ΔLRR)* and *lect-2(null)* mutant animals at L4 stage across 21 minutes. Yellow arrowheads indicate the tips of dendrites growing from the primary dendrite (down). Scale bar, 10 µm. (B) Quantification of the sum of all dendritic growth events in 21 minutes in wild-type, *sax-7(null), dma-1(ΔLRR)* and *lect-2(null)* mutant animals at L4 stage. (C) Quantification of the sum of all dendritic retraction events in 21 minutes in wild-type, *sax-7(null), dma-1(ΔLRR)* and *lect-2(null)* mutant animals at L4 stage. At least 5 animals were imaged for each genotype. n values within each bar. Error bars indicate the standard error of proportion. Statistical comparison was performed using Brown-Forsythe ANOVA with Dunnett’s multiple comparisons test. (D) Time-lapse imaging of developing PVD dendrites (magenta) and SAX-7s::YFP in PLM axons (green) in *Pmec-17::sax-7s::YFP; Pmec-17::mnr-1; ser2prom3::myr-mCherry; sax-7(null)* mutant animals. Representative image of a growing dendrite that was never in contact with the PLM axon within 21 minutes time frame (top). Representative image of a growing dendrite that was always in contact with the PLM axon within 21 minutes time frame (bottom). Yellow arrowheads indicate the tips of dendrites growing from the primary dendrite (down) and PLM axon (above). Scale bar, 10 µm. Schematic images of growing dendrites at 21st minute time point of corresponding conditions (right). PVD dendrites are drawn in magenta and PLM axons containing *Pmec-17::sax-7s::YFP, Pmec-17::mnr-1* are drawn in green. Dashed lines indicate the midline used to quantify the percentage of dendrites that retracted. (E) Quantification of the percentage of dendrites that retracted within 1 hour for dendrites that were in contact with SAX-7, and for dendrites that were not in contact with SAX-7. (F) Quantification of the average length of dendritic retraction per min for both dendrites in contact with SAX-7 and dendrites not in contact with SAX-7. This is calculated as the total length retracted within 21 mins. (G) Quantification of the average length of dendritic growth per min for both dendrites in contact with SAX-7 and dendrites not in contact with SAX-7. This is calculated as the total length that grew within 21 mins. (H) Quantification of the sum of total dendritic growth and retraction per min for both dendrites in contact with SAX-7 and dendrites not in contact with SAX-7. This is calculated as the sum of both total growth and retraction over 21 mins and measures the total movement of the tip of the dendrite. For (E-H), n values within each bar, error bars indicate the standard error of proportion and statistical comparisons were performed using Brown-Forsythe ANOVA with Dunnett’s multiple comparisons test.

To further determine whether ligand binding inhibits continuous growth and retraction, we observed the behavior of dendrites that do or do not contact SAX-7 during dendrite growth. The DMA-1 ligands SAX-7 and MNR-1 are broadly distributed at all levels of PVD branches in wild type animals. Therefore, to observe the behavior of dendrites upon contact with SAX-7 and MNR-1, we spatially restrict the expression of SAX-7 and MNR-1 in mechanosensory PLM neurons, which extends an axonal process that intersects with the secondary branches of PVD. We expressed the ligands SAX-7 and MNR-1 cell-specifically in the mechanosensory neurons (*Pmec-17::sax-7s::YFP; Pmec-17::mnr-1*) in *sax-7(null)* mutant animals (Figure 2D to 2H) and performed time-lapse imaging in L2-L3 animals to record the behavior of secondary dendrites that do or do not contact PLM. As expected, secondary dendrites that are not in contact with PLM exhibit growth and retraction along the dorsal ventral axis similar to *sax-7(null)* mutant animals (Figure 2D). Upon contacting the SAX-7 and MNR-1-expressing PLM, developing dendrites stop growing along the dorsal ventral direction and starts growing along the PLM axon in the anterior-posterior direction (Figure 2D and Smovies). This is consistent with previous studies showing that SAX-7 guides dendritic growth^3^. When we quantified growth and retraction events across dendrites that are contacting or not contacting SAX-7 and MNR-1 expressing PLM, we found that 100% of dendrites not contacting SAX-7 and MNR-1 showed retraction events within an hour, while only a small percentage of dendrites contacting SAX-7 and MNR-1 showed retraction events (Figure 2E). Second, the total retraction length is significantly shorter in dendrites that contact SAX-7 and MNR-1 compared to those that do not make contact (Figure 2F). The growth length was also significantly longer in dendrites without contact, indicating that ligand interaction also inhibits growth of secondary dendrites (Figure 2G). Finally, we summed the total growth and retraction of dendritic tips as indicator of dynamicity (Figure 2H). This measurement quantifies the total movement of the growing dendrites and is also reduced in dendrites that contacted SAX-7 and MNR-1 compared to dendrites that did not contact SAX-7 and MNR-1. Taken together, these results suggest that ligand binding to DMA-1 stabilizes the developing dendrite by preventing continuous dynamicity.

### The LRR of DMA-1 is dispensable for dendrite outgrowth

Because the DMA-1(ΔLRR) can support dynamic dendrite growth, we reason that the growth promoting activity resides in the intracellular domain of DMA-1. We have previously shown that DMA-1 and its co-receptor HPO-30 form a complex to mediate PVD dendrite morphogenesis, and that the cytosolic domains of DMA-1 and HPO-30 directly recruit actin regulators like TIAM-1 and the Wave Regulatory Complex (WRC)^22^. HPO-30 is a claudin-like tetraspanin that is highly enriched in PVD and is necessary for PVD dendrite outgrowth^20,21^. To understand how the cytosolic domains of DMA-1 and HPO-30 promote dendrite growth and branching, we generated a truncated endogenous DMA-1 lacking its intracellular domain (*dma-1(*ΔICD)) and an allele of HPO-30 lacking its C-terminal cytosolic domain (*hpo-30(*ΔICD)). Consistent with previous results, when both the cytosolic tails of DMA-1 and HPO-30 are deleted (*hpo-30(*ΔICD*); dma-1(*ΔICD)), few 4° branches are formed, and overall dendritic branches are dramatically reduced (Figure 3A, 3B, 3C and 3D), indicating that the cytosolic domains of DMA-1 and HPO-30 are required to promote dendritic growth. Deletion of the HPO-30 (*hpo-30(*ΔICD)) alone results in a moderate increase in 4° branches and overall dendritic branches as compared to animals where both cytosolic tails are deleted (*hpo-30(*ΔICD*); dma-1(*ΔICD)) animals (Figure 3A, 3B, 3C and 3D). This indicates that having DMA-1 is sufficient to promote some dendritic growth.

**Figure 3.**
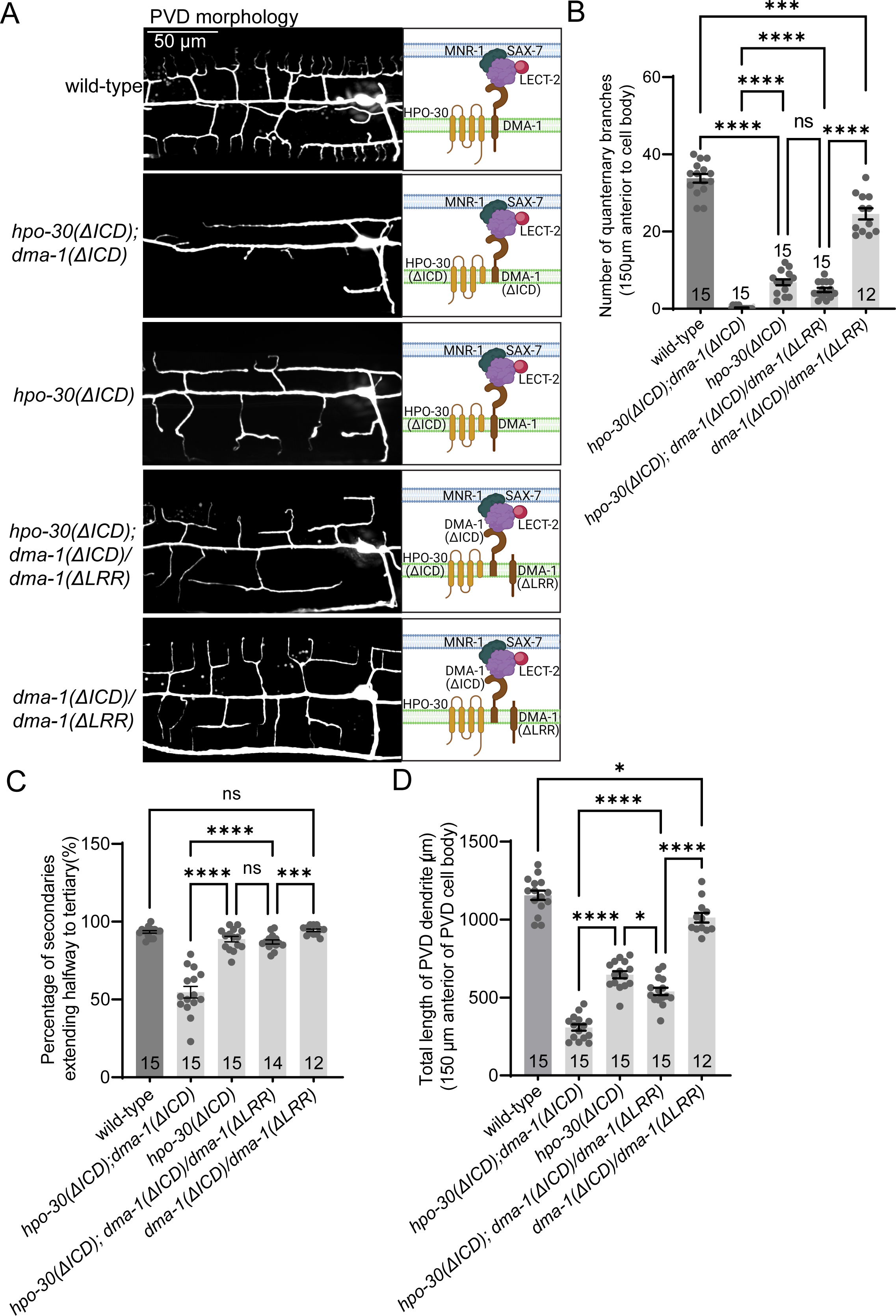
DMA-1 can function as two separate halves. (A) Representative images of PVD dendrite morphology in wild-type, *hpo-30(ΔICD); dma-1(ΔICD), hpo-30(ΔICD), hpo-30(ΔICD); dma-1(ΔICD)/dma-1(ΔICD), dma-1(ΔLRR)/dma-1(ΔLRR)* mutant animals at 1DOA stage. Scale bar, 50 µm. Image representative of 15 animals. Schematic images of molecules that are present in the corresponding animals (left). The cell membrane is drawn in pink, and leaflets of the plasma membrane are differentiated in dark and light colors. MNR-1 is drawn in dark green. SAX-7 is drawn in purple. LECT-2 is drawn in red. DMA-1 is drawn in brown. (B) Quantification of number of quaternary branches in a region 150 µm anterior to the PVD cell body in the indicated genotypes. (C) Quantification of percentage of extended secondaries for the indicated genotypes. Extended secondaries were measured as the percentage of secondary branches per worm that extended at least halfway to the tertiary dendrite line. (D) Quantification of the total length of PVD dendrite, 150 µm anterior to the PVD cell body. For (B-D), n values within each bar, error bars indicate the standard error of proportion. Statistical comparison was performed using Brown-Forsythe ANOVA with Dunnett’s multiple comparisons test.

Based on our results so far, we hypothesized that the intracellular domain of DMA-1 promotes dendrite growth while the extracellular domain inhibits growth through ligand binding. One important question is to understand how these two functions are coordinated for proper dendritic development. To answer this, we expressed DMA-1 as separate halves. If DMA-1 extracellular and intracellular domains indeed have distinct functions, expressing one copy of extracellular, and one copy of intracellular DMA-1 would reconstitute DMA-1 and function to promote dendritic growth. We analyzed heterozygous *hpo-30(ΔICD); dma-1(ΔICD)/ dma-1(ΔLRR)* animals generated by crossing *hpo-30(ΔICD); dma-1(ΔICD)* animals with *hpo-30(ΔICD); dma-1(ΔLRR)* animals. In these animals, HPO-30 lacks its cytosolic tail and DMA-1 exists as two separate proteins, one containing the extracellular domain and transmembrane domain (DMA-1(ΔICD)) and the other containing the transmembrane domain and cytoplasmic domain (DMA-1(ΔLRR)). Remarkably, we observed no significant difference in the number of 4° branches, and the length of 2° branches in these animals compared to *hpo-30(ΔICD)* animals (Figure 3A, 3B and 3C). The total dendrite length of the *hpo-30(ΔICD); dma-1(ΔICD)/ dma-1(ΔLRR)* animals are slightly shorter than that of the *hpo-30(ΔICD)* animals, but longer than the *hpo-30(ΔICD); dma-1(ΔICD)* animals (Figure 3D). This might be due to differences in the number of *dma-1* expressed. Importantly, both *hpo-30(ΔICD); dma-1(ΔICD)/ dma-1(ΔLRR)* and *dma-1(ΔICD)/ dma-1(ΔLRR)* animals show ordered dendritic menorahs (Figure 3A). These results argue strongly that the functions of extracellular and intracellular domains of DMA-1 are distinct, and these functions can be achieved as separate proteins.

As cell surface receptors are presumed to receive extracellular cues and activate intracellular signals, these results are surprising and support two notions. First, the cytoplasmic domain of DMA-1 promotes dendrite growth and branching in the absence of its extracellular domain. Second, the function DMA-1 receptor can be reconstituted as separate halves of DMA-1, each with distinct DMA-1 function. The extracellular domain is required to guide the growing dendrite for the formation of the proper shape and morphology of dendrites while the intracellular domain promotes dendritic growth. From these results, we propose that ligand binding through the extracellular domain of receptor DMA-1 is not required for the intracellular domain to activate downstream signaling pathways to drive cytoskeletal rearrangement. Instead, ligand binding stabilizes dendrites and inhibits continuous dynamicity.

### Diffusible ligand-free DMA-1 is responsible for promoting dendritic growth

The fact that extracellular and intracellular functions of DMA-1 can be reconstituted as separate proteins strongly suggests that there are two populations of DMA-1 *in vivo*. To examine DMA-1 subcellular localization during dendritic development, we visualized animals with endogenous DMA-1::GFP that was generated previously by CRISPR-Cas9^29^. We previously reported two pools of DMA-1 with distinct dynamics: a diffuse pool that is mobile, outlines the PVD cell body and present along the dendrites, and a punctate pool that can be seen both in the soma and along the dendrites^29^ (Figure 4A). It was also suggested in the same study that diffuse signals present near the cell body are freely moving DMA-1 on the cell membrane, while the bright puncta are DMA-1 trapped within vesicles^29^. Consistent with previous findings, when we performed fluorescence recovery after photobleaching (FRAP) experiment on both diffuse and punctate dendritic DMA-1::GFP, we observed that the diffuse signal recovered to about 50% of the pre-bleaching level within 45 seconds, indicating the presence of a pool of diffusible DMA-1 on the plasma membrane (Figure 4B). In contrast, the puncta show little recovery after photobleaching, consistent with DMA-1 being in internal vesicles with discontinuous membranes (Figure 4B).

**Figure 4.**
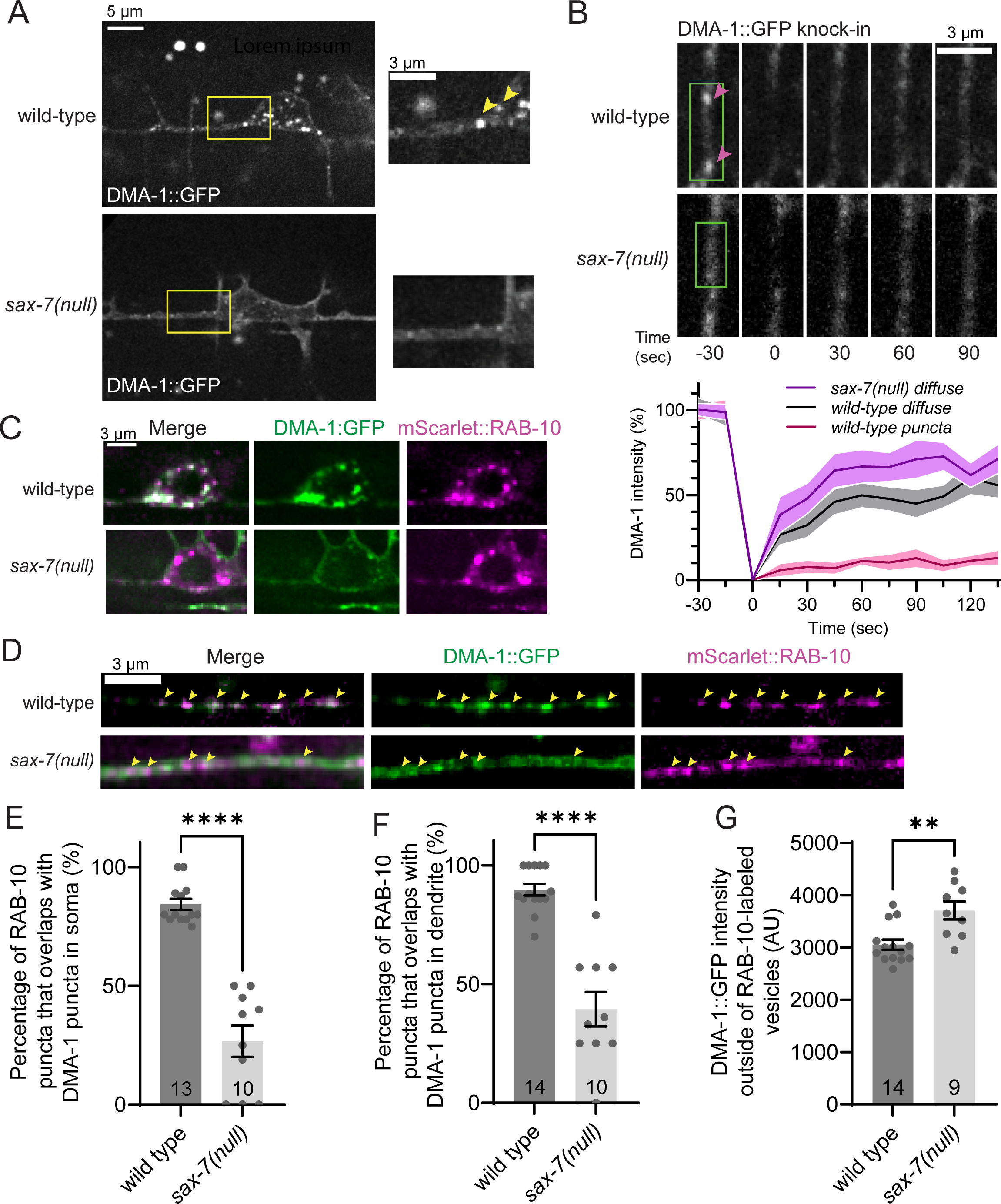
SAX-7 is required for the endosomal localization of DMA-1. (A) Endogenous DMA-1::GFP knock-in in wild-type and *sax-7(null)* mutants. Yellow boxes highlight the locations of magnified views on the right. Arrowheads indicate punctate structures representing putative endosomes. (B) Fluorescence recovery after photobleaching in wild-type and *sax-7(null)* mutants expressing endogenous DMA-1::GFP. The green rectangle indicates the region of primary dendrite that was bleached at time = 0 sec. Arrowheads indicate DMA-1::GFP puncta. Error bars indicate the standard error of proportion from independent animals (n = 36 diffuse regions analyzed from 14 individual wild-type worms; n = 20 puncta analyzed from 9 individual wild-type worms; n= 28 diffuse regions from 15 *sax-7(null)* individual animals). (C-D) Colabeling of endogenous DMA-1::GFP with endogenous mScarlet:RAB-10 (mScarlet::RAB-10; expressed cell-specifically in PVD) in the soma (C) and primary dendrite (D) in the *sax-7(null)*. Arrowheads indicate locations of mScarlet::RAB-10 puncta. (E-F) Quantification of the percentage of mScarlet::RAB-10 puncta that overlaps with DMA-1:GFP puncta in the soma (E) and primary dendrite (F) in the *sax-7(null)*. (G) Quantification of DMA-1::GFP fluorescence intensity measured outside of mScarlet:RAB-10 puncta in the primary dendrite in *sax-7(null)*. For (E-G), n values within each bar, error bars indicate the standard error of proportion. Statistical comparison was performed using a Mann-Whitney nonparametric t-test in (E-G).

Then, we investigated the identity of these DMA-1-containing internal vesicles. As RAB-10, a small GTPase localized to recycling endosomes was previously shown to be essential for DMA-1 trafficking and PVD dendrite arborization^30,31^, we asked if DMA-1 is present in RAB-10-labeled vesicles. To visualize the localization of endogenous RAB-10 vesicles in PVD dendrites, we used the previously described flippase (FLP)-mediated labeling strategy^32^ to generate cell-specific, endogenously-tagged mScarlet::RAB-10. We found that RAB-10 localizes to puncta distributed throughout the PVD soma and dendrites (Figure 4C and 4D). In wild-type worms, we observed a high degree of colocalization between DMA-1 puncta and RAB-10 labeled vesicles in both the soma (Figure 4C and 4E) and dendrites (Figure 4D and 4F). Thus, the DMA-1 puncta observed in wild-type worms likely represent an endosomal pool of DMA-1 that is largely localized to RAB-10 positive recycling endosomes. Together with our earlier observations, these data suggest that there are two populations of DMA-1, one population that is relatively diffusible and present on the cell surface and another population that is traveling through endosomes. Many membrane receptors undergo ligand-induced endocytosis, which can mediate the strength or duration of intracellular signaling processes by physically reducing the concentration of cell surface receptors available to the ligand^33^. We asked whether the interaction between DMA-1 and its ligand SAX-7 induces its endocytosis and influence its subcellular distribution. To understand the relationship between the plasma membrane and endosomal DMA-1, we examined DMA-1::GFP in *sax-7(null)* mutants. We observed that *sax-7(null)* mutants show fewer DMA-1 puncta in both the cell soma and along the dendrites, while the diffuse DMA-1 on the cell membrane remains intact (Figure 4A). Furthermore, when we performed FRAP experiments on the diffusible pool of DMA-1 in *sax-7(null)* mutants, the rate of recovery of DMA-1::GFP was faster than wild-type animals (Figure 4B). The fluorescence recovery was also more complete in *sax-7(null)* mutants. To further illustrate this, we expressed mScarlet::RAB-10 in *sax-7(null)* mutant animals and quantified the percentage of DMA-1::GFP that overlaps with mScarlet::RAB-10, as well as the intensity of DMA-1::GFP outside of RAB-10. We observed that the percentage of DMA-1::GFP that co-localizes with mScarlet::RAB-10 both in the soma (Figure 4C and 4E) and dendrites (Figure 4D and 4F) was drastically reduced compared to wild-type animals. In addition, there is an increase in DMA-1::GFP intensity outside of RAB-10 (Figure 4G), indicating that there is a larger portion of diffusible DMA-1. Together, these data suggest that SAX-7 and possibly ligand binding induces DMA-1 internalization into the RAB-10 endosomes.

### KPC-1 cleaves HPO-30

Because SAX-7 and ligand binding is not required for DMA-1 mediated dendrite growth, we reason that there must be other mechanisms to activate DMA-1 for dendrite outgrowth. To uncover these mechanisms, we examined known mutants displaying strong PVD dendrite defects. Previous studies showed that the proprotein convertase KPC-1 is required for PVD dendrite growth (Figure S1A) and that *kpc-1(null)* mutants have increased DMA-1 levels^24–26^. However, it is unclear how furin-like KPC-1 regulates dendrite morphogenesis through DMA-1. To identify additional KPC-1 interactors that promote dendrite growth, we performed a genetic modifier screen on a weak *kpc-1* allele, *kpc-1(xr58).* While *kpc-1(null)* mutants have dramatically reduced secondary branches and no quaternary branches, *kpc-1(xr58)* mutants have a mild reduction of secondary branches and dramatically reduced quaternary branches (Figure S1A, S1C and S1D). Our genetic modifier screen sought additional mutations that enhance the dendrite phenotype of *kpc-1(xr58)*. From this screen, we identified *wy1008* (*hpo-30(E100K)*), which is a point mutation in HPO-30 that causes an E100K amino acid change in the first extracellular loop of HPO-30 (Figure S1B). *kpc-1(xr58)*; *hpo-30(E100K)* double mutants showed a strong loss of dendrite similar to the *kpc-1(null)* mutants (Figure S1A, S1B, S1C and S1D). *hpo-30(E100K)* is likely a partial loss-of-function allele since the *hpo-30(E100K)* dendrite defects are milder than those of *hpo-30(null)* mutants, which generally fail to form quaternary branches but have longer secondary branches compared to the *kpc-1(null)* mutants (Figure S1B, S1C, and S1D). Together, these data indicate a genetic interaction and possibly molecular interaction between *kpc-1* and *hpo-30* to regulate dendrite outgrowth.

Since KPC-1 is predicted to be a proprotein convertase homologous to furin, we asked whether KPC-1 cleaves HPO-30 in PVD neurons. To test this, we generated worms expressing HPO-30 fused with GFP on the C-terminus (HPO-30::GFP) under the control of a PVD promoter (*ser2prom3::hpo-30::GFP*). We then asked whether HPO-30 exists in shorter cleaved fragments in worm lysates by performing a Western blot using a GFP antibody. Indeed, in wild-type worms expressing HPO-30::GFP, we detected both full-length HPO-30::GFP as well as a smaller fragment corresponding to a C-terminal cleavage fragment (Figure 5A). Moreover, the shorter C-terminal fragment was not observed in *kpc-1(null)* mutants (Figure 5A), indicating that HPO-30 cleavage is dependent on KPC-1. As HPO-30 is the co-receptor for DMA-1, we tested if the DMA-1 is necessary for the cleavage of HPO-30. We observed that HPO-30::GFP cleavage is not affected in *dma-1(null)* mutants, suggesting that HPO-30 cleavage does not require DMA-1 (Figure 5A). To further verify that furin-like proteases can cleave HPO-30, we expressed either C-terminally-tagged HPO-30::GFP or N-terminally-tagged FLAG::HPO-30 in *Drosophila* S2 cells and treated the cells with a furin inhibitor (decanoyl-RVKR-CMK). We then performed Western blots using anti-GFP to detect cleaved C-terminal HPO-30 fragments or anti-FLAG antibodies to detect cleaved N-terminal HPO-30 fragments and observed that cleaved HPO-30 was greatly reduced in both cases (Figure 5B). This suggests that KPC-1 likely cleaves HPO-30 in both *C. elegans* and Drosophila cells.

**Figure 5.**
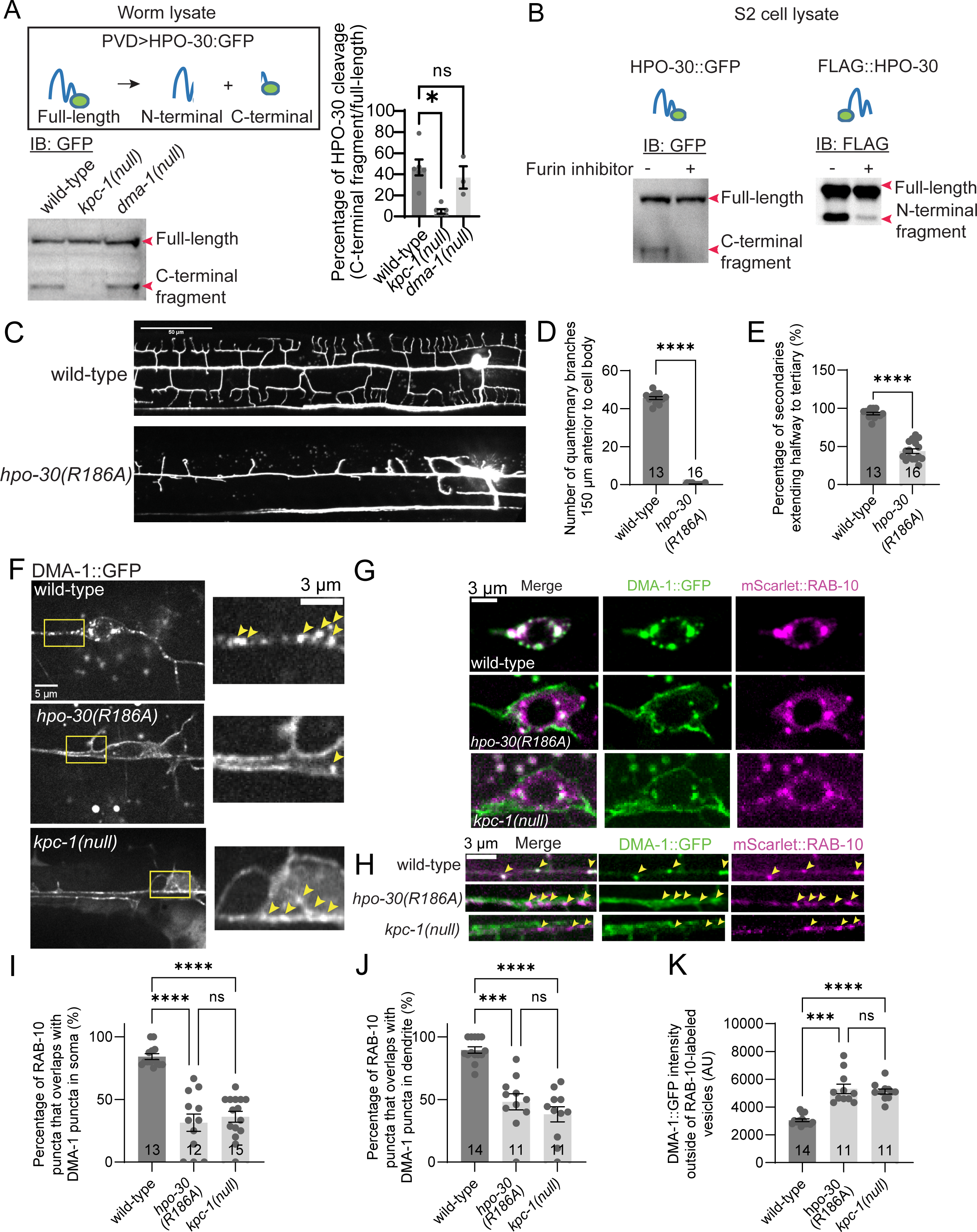
Cleavage of HPO-30 by KPC-1 facilitates DMA-1 recycling and is necessary for dendrite outgrowth. (A) Western blot against GFP in worms expressing an HPO-30::GFP transgene expressed under a PVD promoter. Top: Schematic of full-length HPO-30::GFP protein and putative N-terminal and C-terminal cleavage fragments. Bottom left: Western blot using anti-GFP antibody on whole-worm lysate of wild-type, *kpc-1(null)*, and *dma-1(null)* worms. Right: Quantification of HPO-30 cleavage, calculated as intensity of C-terminal fragment out of total intensity. Statistical comparison was performed using Brown-Forsythe ANOVA with Dunnett’s multiple comparisons test (n = 6 blots for wild type and *kpc-1(null)*; n=3 blots for *dma-1(null)*). (B) Western blot against GFP in cell lysates from Drosophila S2 cells expressing HPO-30::GFP (left) or FLAG::HPO-30 (right) constructs. Cells were grown in the absence (-) or presence (+) of the furin inhibitor decanoyl-RVKR-CMK. (C) Representative images of PVD morphology in wild-type and *hpo-30(R186A*) mutants at the L4 stage. *hpo-30(R186A)* mutants were generated by CRISPR into the endogenous *hpo-30* locus. (D) Quantification of number of quaternary branches in a region 150 µm anterior to the PVD cell body for *hpo-30(R186A)*. (E) Quantification of percentage of extended secondaries in *hpo-30(R186A)*. In (D and E), n values within each bar, error bars indicate the standard error of proportion. Statistical comparison was performed using Brown-Forsythe ANOVA with Dunnett’s multiple comparisons test. (F) Endogenous DMA-1::GFP in wild-type, *hpo-30(R186A)* and *kpc-1(null)*. Yellow rectangles indicate locations of magnified views on the right. Arrowheads indicate punctate structures representing putative endosomes. (G and H) Colabeling of endogenous DMA-1::GFP with endogenous mScarlet::RAB-10 (mScarlet::RAB-10; expressed cell-specifically in PVD) in the cell bodies (G) and primary dendrite (H) of wild-type, *hpo-30(R186A)* and *kpc-1(null)* animals. Arrowheads indicate locations of mScarlet::RAB-10 puncta. (I and J) Quantification of the percent of mScarlet::RAB-10 puncta that overlaps with DMA-1::GFP puncta in the cell bodies (I) and primary dendrites (J). (K) Quantification of DMA-1::GFP fluorescence intensity measured outside of mScarlet::RAB-10 puncta. For (I, J and K), n values within each bar, error bars indicate the standard error of proportion. Statistical comparison was performed using Brown-Forsythe ANOVA with Dunnett’s multiple comparisons test.

### HPO-30 cleavage by KPC-1 is required for dendritic growth and regulates DMA-1 internalization to the recycling endosome

To identify the cleavage site in HPO-30, we scanned the HPO-30 protein sequence for the furin recognition consensus motif “RXXR”^34^ and individually mutated these arginines to alanines (R to A). We systematically expressed these R to A constructs in S2 cells and assayed them for HPO-30::GFP cleavage by western blot. We found that either a R186A or R189A mutation eliminated the cleavage product (Figure S2A). Consistent with the canonical furin cleavage motif, these two residues are part of an “RRER” sequence (residues 186-189) in the second extracellular loop of HPO-30 (Figure S2A), which is predicted to reside in the lumen of secretory space. The location of this cleavage site is also consistent with the size of the cleaved HPO-30 fragments.

Next, we asked if HPO-30 cleavage is required for PVD dendrite morphogenesis. We first generated animals containing the R186A mutation in HPO-30 using CRISPR/Cas9. We found that *hpo-30(R186A)* mutants showed a dramatic loss of dendrites similar to *hpo-30(null)* (Figure 5C, 5D, 5E and S1B). Quaternary branches were completely absent (Figure 5C and 5D), while short secondary branches remained at about 50% of the level of wild-type controls (Figure 5C and 5E). These results indicate that HPO-30 cleavage is required for its function in promoting dendritic growth.

As HPO-30 forms a complex with DMA-1 to promote dendritic growth, we sought to understand how HPO-30 cleavage activates this receptor complex. First, we examined if blocking HPO-30 cleavage prevents HPO-30 from binding to DMA-1 by performing a co-immunoprecipitation experiment in *Drosophila* S2 cells. We expressed DMA-1 tagged with HA along with either wildtype or R186A HPO-30 tagged with GFP, then immunoprecipitated HPO-30 with an anti-GFP antibody, and probed DMA-1 using an anti-HA antibody. We observed that uncleavable HPO-30(R186A) can still bind to DMA-1 (Figure S2B), suggesting that the HPO-30/DMA-1 complex is still intact and does not require HPO-30 cleavage.

We then asked whether HPO-30 cleavage affects DMA-1 localization. To do so, we examined DMA-1::GFP in *hpo-30(R186A)* and *kpc-1(null)* mutants where HPO-30 cleavage does not occur. Remarkably, punctate internal DMA-1::GFP signals were reduced in the soma and along the dendrite, but the plasma membrane signal was increased in both *hpo-30(R186A)* and *kpc-1(null)* animals (Figure 5F). Consistent with this, when we looked at the co-localization of DMA-1::GFP and mScarlet::RAB-10, we observed that there was a reduction in the percentage of DMA-1::GFP signals that overlapped with mScarlet::RAB-10 in the soma (Figure 5G and 5I) and along the dendrite (Figure 5H and 5J) in both *hpo-30(R186A)* and *kpc-1(null)* animals. In addition, there was an increase in diffuse DMA-1 signal in both *hpo-30(R186A)* and *kpc-1(null)* animals (Figure 5K). These data suggest that HPO-30 cleavage by KPC-1 promotes the internalization of DMA-1 into endosomes. Interestingly, when we visualized HPO-30::GFP in *kpc-1(null)* mutants, we also observed that HPO-30::GFP becomes more diffuse with fewer intracellular HPO-30 puncta compared to wild type animals, similar to DMA-1::GFP (Figure S2C). These data are consistent with a model that KPC-1 cleaves HPO-30 and induces the endocytosis of DMA-1/HPO-30 complex into recycling endosomes, which is essential for the growth promoting functions of DMA-1 and HPO-30.

### Endosomal DMA-1 is important for PVD dendrite development

Through a separate genetic screen, we identified a novel allele of *dma-1*, *zac98* (*dma-1(C470Y)*) that contains a single C470Y point mutation in the extracellular domain of DMA-1. *dma-1(C470Y)* mutant animals displayed dramatic dendrite defects that are indistinguishable from the *kpc-1(null)* mutants but surprisingly distinct from *dma-1(null)* (Figure S3A, S3B and S3C), suggesting that the C470Y mutation does not simply cause loss of *dma-1* function. When we assessed the localization of DMA-1(C470Y)::GFP in *dma-1(C470Y)* mutant animals, DMA-1(C470Y)::GFP exhibited increased plasma membrane localization and reduced internal puncta in the soma and the dendrites compared to the wild-type (Figure S3D), similar to *kpc-1(null)* and *hpo-30(R186A)* mutants (Figure 5E to 5J).

To further understand how the regulation of the endosomal and plasma membrane pools of DMA-1 affect dendrite development, we looked at the co-localization of DMA-1::GFP and mScarlet::RAB-10 in *dma-1(C470Y)* mutants. We observed a reduction in the percentage of DMA-1::GFP signal that overlapped with mScarlet::RAB-10 in the soma and along the dendrite in *dma-1(C470Y)* animals (Figure S3E to S3H). Finally, there is an increase in diffuse DMA-1 in *dma-1(C470Y)* animals (Figure S3I). These data suggest that DMA-1 needs to be internalized into endosomes for the proper formation of the PVD neuron.

To further dissect the function of DMA-1 internalization in the formation of the PVD, we wondered if the endocytic process is required to dissociate ligand-bound DMA-1 from its transmembrane ligand SAX-7 while it moves into endosomes to then be recycled onto the membrane as ligand-free DMA-1 to promote growth activity. To test this hypothesis, we performed FRAP analyses to measure the proportion of mobile DMA-1::GFP in mutants (*kpc-1(null)*, *hpo-30(R186A)* and *dma-1(C470Y)*) with defects in DMA-1 endocytosis, resulting in largely mis-localized DMA-1 on the plasma membrane. If DMA-1 endocytosis is required to form ligand-free DMA-1 on plasma membrane, mutants with defects in DMA-1 endocytosis would have a reduction of mobile DMA-1 since we have shown earlier than ligand-free DMA-1 is more mobile than ligand-bound DMA-1 (Figure 4B). Indeed, we saw that DMA-1::GFP recovered poorly in these mutants after photobleaching, suggesting that mobile DMA-1 is reduced in these mutants as compared to wild-type (Figure 6A and 6B). Because these mutants showed increased DMA-1 plasma membrane localization where DMA-1 interacts with its ligands, we wondered if DMA-1 was immobilized through ligand binding. To test this, we removed the ligand SAX-7 in *kpc-1(null)*, *hpo-30(R186A)* and *dma-1(C470Y)* mutants by creating double mutants between *sax-7(null)* and each of the mutants. We then measured the proportion of mobile DMA-1 using FRAP. Strikingly, DMA-1::GFP recovers significantly faster and more completely in the double mutants (*kpc-1(null); sax-7(null)*, *hpo-30(R186A); sax-7(null)* and *dma-1(C470Y); sax-7(null))* as compared to the single mutants. This observation strongly indicates that ligand binding immobilizes DMA-1 in these three mutant backgrounds, resulting in PVD defects (Figure 6A, 6C, 6D and 6E). Moreover, the three double mutants which have an increased proportion of diffused DMA-1 exhibit longer dendritic branches than the corresponding single mutants (Figure S4A and S4B), suggesting that the binding between SAX-7 and DMA-1 immobilizes the receptor and inhibits branch growth. Together, these data are consistent with the two notions. First, the ligand-free DMA-1 present on the surface is generated by endosomal recycling and is required to promote dendritic growth. Second, while ligand binding inhibits growth and prevents retraction to achieve selective stabilization, it is also required for DMA-1 recycling to generate mobile ligand-free DMA-1 on the cell membrane necessary for dendritic growth.

**Figure 6.**
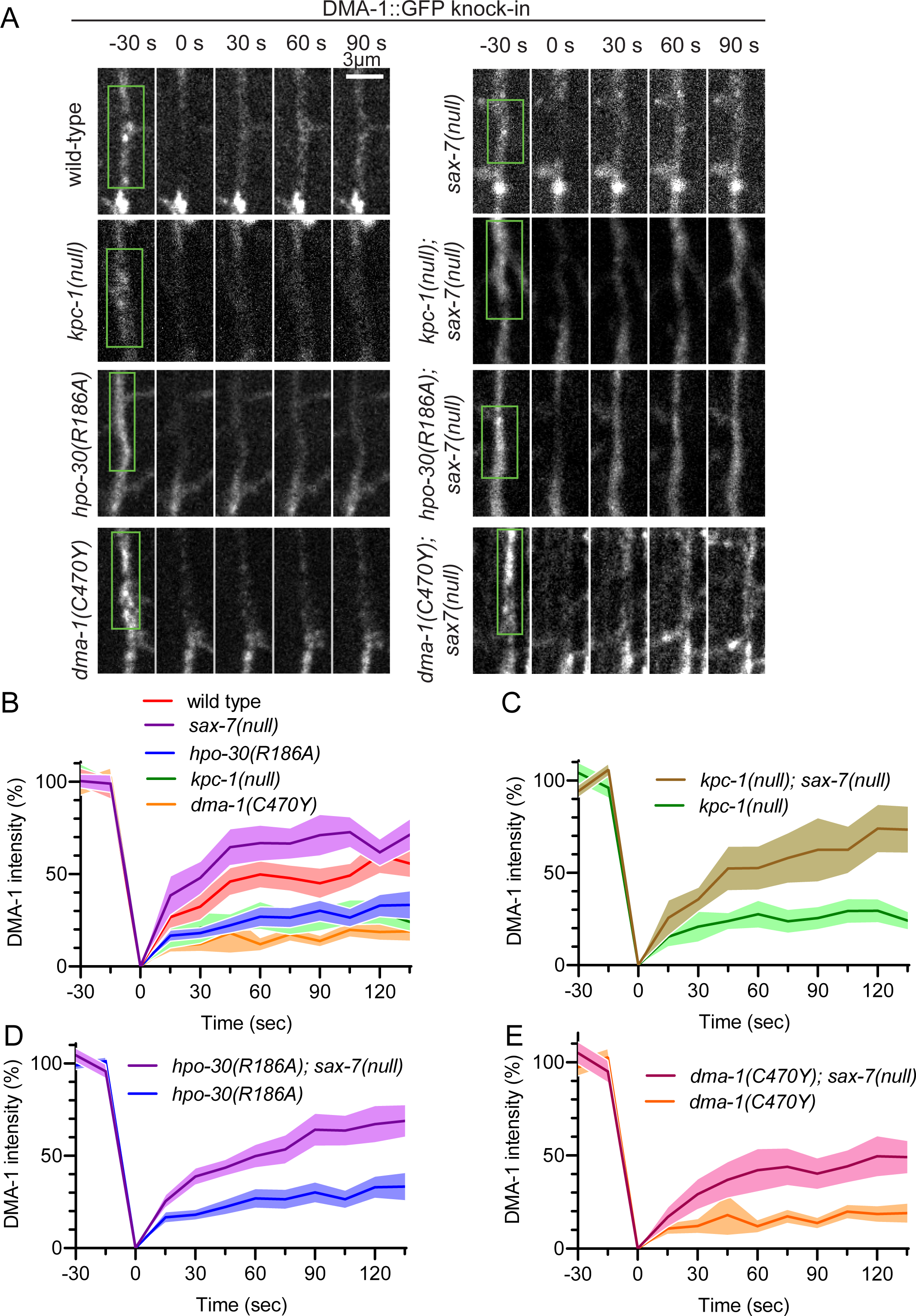
Endocytosis/recycling of DMA-1 provides a diffusible pool of DMA-1. (A) Representative images of fluorescence recovery after photobleaching in the indicated genotypes expressing endogenous DMA-1::GFP. Green rectangle indicates the region of primary dendrite that was bleached at time = 0 s. Scale bar = 3µm. (B-E) Fluorescence recovery after photobleaching analysis of endogenous DMA-1::GFP in the indicated genotypes showing in (A). For (B-E), error bars indicate 95% confidence interval. Wild-type: 36 diffuse regions from 14 animals, *kpc-1(null)*: 26 diffuse regions from 12 animals, *kpc-1(null); sax-7(null)*: 20 diffuse regions from 10 animals, *hpo-30(R186A)*: 23 diffuse regions from 12 animals, *hpo-30(R186A); sax-7(null)*: 28 diffuse regions from 15 animals, *dma-1(C470Y)*: 20 diffuse regions from 10 animals, *dma-1(C470Y); sax-7(null):* 28 diffuse regions from 14 animals.

### DMA-1 recycling promotes DMA-1 diffusion into nascent dendrites

Since DMA-1 functions as guidance receptor, why does enhanced localization of DMA-1 on the plasma membrane and reduced endosomal localization lead to near-complete failure of dendrite morphogenesis? To further probe this question, we performed time-lapse imaging experiments to pinpoint the developmental deficits that led to the dramatic loss of dendrites in mutant animals with defects in DMA-1 recycling (*kpc-1(null)*, *hpo-30(R186A)* and *dma-1(C470Y)*). In wild-type animals, dendritic filopodia often undergo rounds of growth and retraction, which ultimately results in net growth (Figure 7A, 7B and 7C). In contrast, we found that in *kpc-1(null)*, *hpo-30(R186A)* and *dma-1(C470Y)* mutants with depleted endosomal DMA-1, short filopodia initiated normally but failed to extend beyond 1.5µm in length. These short filopodia retracted within a few minutes, resulting an overall slower growth speed (Figure 7A, 7B and 7C). This further suggests that the reduction in mobile ligand-free DMA-1 impedes dendritic growth.

**Figure 7.**
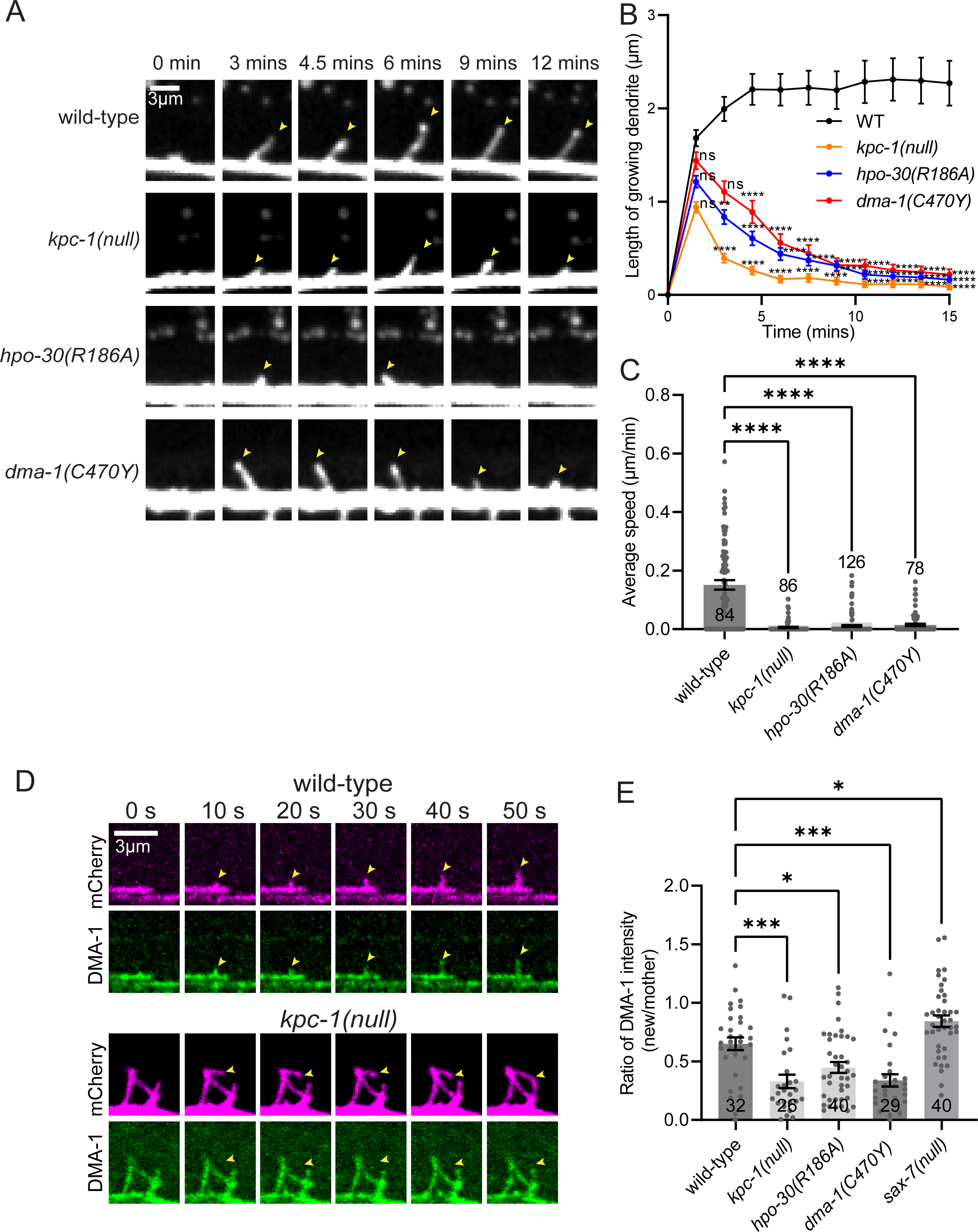
Endosomal DMA-1 is required for rapid recruitment of DMA-1 to newly formed dendrites. (A) Time-lapse imaging of developing PVD secondary dendrites in wild-type, *kpc-1(null)*, hpo*-30(R186A)*, and *dma-1(C470Y)* animals. Yellow arrowheads indicate growth and retraction events of the dendritic tips. (B) A plot of growing secondary dendrites length over time in the indicated genotypes. Number of growing secondary dendrites used in: wild-type = 84, *kpc-1(null)* = 86, *hpo-30(R186A)* = 126 and *dma-1(C470Y)* = 78. Error bars indicate the standard error of proportion. Statistical comparison was performed using Brown-Forsythe ANOVA with Dunnett’s multiple comparisons test. (C) Quantification of the average speed of secondary dendrite growth in wild-type, *kpc-1(null)*, hpo*-30(R186A)*, and *dma-1(C470Y)* animals. n values within each bar and more than 4 independent animals were imaged for each genotype. Error bars indicate the standard error of proportion. (D) Time-lapse imaging of developing PVD dendrites with DMA-1::GFP (green) and mCherry (magenta) in wild-type and *kpc-1(null)* animals. (E) Quantification of the ratio of DMA-1::GFP intensity (newly formed dendrite/mother dendrite) in wild-type, *kpc-1(null)*, hpo*-30(R186A)*, and *dma-1(C470Y)* animals. More than 13 independent animals were imaged for each genotype. For (B, C and E), statistical comparison was performed using Brown-Forsythe ANOVA with Dunnett’s multiple comparisons test.

Next, we wondered if this reduction in overall dendritic growth speed is due to the lack of DMA-1 in the newly generated filopodia. To characterize the presence of DMA-1 in nascent filopodia, we performed time-lapse imaging in animals expressing endogenous DMA-1::GFP and a PVD marker (*ser2prom3::mCherry*), and asked how quickly DMA-1::GFP enters a newly formed filopodia. This is quantified by calculating the ratio between the intensity in nascent filopodia and the intensity in the mother branch at 7.5 seconds after filopodia formation. In wild-type animals, DMA-1::GFP is enriched in the nascent filopodia within seconds after filopodia growth (Figure 7D). However, in *kpc-1(null), hpo-30(R186A)* and *dma-1(C470Y)* mutants, it took significantly longer for DMA-1::GFP to appear and be enriched in the nascent filopodia (Figure 7D and S5). *kpc-1(null)*, *hpo-30(R186A)* and *dma-1(C470Y)* mutants showed reduced enrichment of DMA-1 on nascent branches as compared to wild-type animals (Figure 7D, 7E and S5). These data suggest that while the DMA-1 recycling mutants have higher levels of DMA-1 on their plasma membrane, DMA-1 is not rapidly recruited to nascent dendritic filopodia. Without DMA-1 in the nascent filopodia, the branches retract, leading to a lack of dendritic arbors. Our data suggest that freely diffusible, ligand-free DMA-1 can laterally diffuse (Figure 7D and 7E) into nascent filopodia to facilitate dendrite growth. However, because of the striking colocalization between punctate DMA-1 and recycling endosomes, it is possible that exocytosis of DMA-1-containing recycling endosomes contributes to the enrichment of DMA-1 at the tips of the nascent filopodia. To test this idea, we used endogenously labeled GFP::RAB-10 to track the localization and dynamics of RAB-10-positive endosomes during filopodia formation. We found that RAB-10 puncta were distributed sparsely along the dendrite, with no obvious enrichment at the sites of filopodia formation (Figure S6). In most of *de novo* filopodia formation events, there was no obvious RAB-10 positive puncta localized near or inside of the nascent filopodia, suggesting that nascent filopodia formation does not require local RAB-10 endosomal exocytosis.

Together, these data reveal a new model for how ligand receptor systems mediate stochastic dendrite growth to build stereotyped dendritic arbors (Figure S7). First, SAX-7 interacts with DMA-1 on the plasma membrane to stabilize dendritic branches and build dendritic arbors based on SAX-7 localization. Second, SAX-7 also induces DMA-1 endocytosis and drive DMA-1 into the RAB-10 endosomes. Through this process, DMA-1 dissociates from the transmembrane SAX-7 and becomes ligand-free. Third, once the receptor is recycled on the plasma membrane, the ligand-free DMA-1 diffuses into nascent filopodia and promotes dynamic, stochastic growth. Once DMA-1 binds to new SAX-7, the cycle repeats. This new model provides a molecular mechanism to integrate the stochastic growth during dendritic morphogenesis and the stereotyped dendritic arbor. Ligand-free receptor promotes stochastic, dynamic growth, while ligand-bound receptors stabilize dendrites and inhibit dynamic growth to achieve the arbor shape.

## Discussion

During dendritic morphogenesis, intrinsic growth molecules and extrinsic guidance cues act in concert to determine the proper size and shape of a neuron. Altered dendritic morphology is associated with neuronal diseases such as autism spectrum disorders^35^ and Alzheimer’s disease^36^. Here, we reveal mechanisms of dendritic morphogenesis that not only consolidate our current knowledge, but further our understanding of how dendritic arborization is regulated at the molecular level. Our data suggest that DMA-1 functions as an adhesion molecule to stabilize growing dendrites through its extracellular ligand-binding domain, but also promotes dendritic growth through its cytoplasmic domain. We show that this growth promoting function is dependent on KPC-1 cleavage of HPO-30 which activates the DMA-1/HPO-30 receptor complex. We demonstrate that a pool of freely diffusible DMA-1 needs to be generated by recycling endosomes, which then rapidly diffuse into nascent filopodia to promote dendritic growth. Together, our data uncover novel aspects of the cell biology of guidance receptors and further our understanding of neuronal development.

Dynamic imaging of dendrite branching in Drosophila neurons has shown that dendrite tips exhibit stochastic growth, pause and retraction^10,14,26,37^, and computational modeling suggests that this dynamic nature and the rapid transitions between growth and retraction shape and maintain dendritic morphology^38^. In *C. elegans*, the developing PVD dendrites also exhibit continuous growth and retraction^14,26,39^. Our results surprisingly show that while ligand binding to DMA-1 is required for guidance, it also inhibits dendrite growth instead of promoting growth. Therefore, we propose that the stereotypical shape of the PVD neuron’s dendritic arbor is first achieved through stochastic intrinsic growth mediated by ligand-free DMA-1 and not by the binding of DMA-1 to its ligand. This is then followed by selective growth inhibition and branch stabilization through ligand-bound DMA-1. This model challenges the conventional model in neuronal morphogenesis where ligand binding to cell surface receptors induces a conformational change in the cytoplasmic tail to drive axon and dendrite growth. Moreover, our results suggest that ligand-free DMA-1 is diffusible, and that increasing the proportion of ligand-free DMA-1 promotes dendritic growth.

Previous studies have shown that DMA-1 is localized on the cell membrane and in intracellular vesicles^1,21,31^, and that RAB-10 is required for the recycling of DMA-1 onto the plasma membrane to function as a guidance receptor^30,31^. Our results, however, suggest that receptor localization to the plasma membrane alone is not sufficient for its function. Instead, the recycling of the receptor from the plasma membrane and into endosomes is also important for dendritic development. We propose the internalization of guidance receptors generates a diffusible pool of plasma membrane-localized receptors that is essential for dendritic filopodia outgrowth.

Given the notion that HPO-30 and DMA-1 form a receptor complex, our data indicate that KPC-1 cleavage of HPO-30 activates the receptor complex to promote dendritic morphogenesis. Because KPC-1 functions cell-autonomously in PVD^24,26^ and DMA-1 is not required for HPO-30 cleavage by KPC-1 (Figure 5A) to promote dendrite formation, it is plausible that KPC-1 cleavage is independent of the formation of DMA-1/HPO-30 complex and ligand binding to the receptor. Interestingly, we found increased membrane DMA-1::GFP and reduced endosomal DMA-1::GFP in the non-cleavable *hpo-30(R186A)* mutant allele (Figure 5F and 5G). These data indicate that the full length HPO-30 prevents internalization, but this effect is negated once KPC-1 cleaves HPO-30. DMA-1 binds to both the cleavable and non-cleavable HPO-30 (Fig. S2B). We speculate that when DMA-1 binds to full-length HPO-30, its lateral diffusion and endocytosis are inhibited, possibly due to claudin’s ability to form cis interactions with itself^40^. When DMA-1 binds to cleaved HPO-30, it is then able to be endocytosed and recycled back to the plasma membrane, providing a ligand-free receptor that can diffuse into newly formed filopodia. HPO-30 has also been found to maintain surface levels of levamisole-dependent AChRs at neuromuscular junctions^40^, suggesting the regulation of surface receptor levels may be a generalizable function of HPO-30 and perhaps other claudins.

Like many sensory receptors on the cell surface, guidance receptors undergo endocytosis and recycling. The biological functions of membrane trafficking of receptors are three-fold. First, receptor endocytosis provides a means to dynamically regulate the level of guidance receptors on the plasma membrane. For example, Semaphorin binding to Plexin leads to internalization of Plexin and reduced sensitivity to the repulsive Semaphorin gradient^15,41^. In addition, Eph receptors are endocytosed upon ligand binding and receptor activation, and subsequently sorted into late endosomes and lysosomes for degradation^33^. In this study, we have also discovered that the binding of SAX-7/L1CAM to DMA-1 is necessary for the internalization of DMA-1 into recycling endosomes.

Second, receptor endocytosis generates endosomes that can serve as signaling hubs. The importance of signaling endosomes has been extensively described for a variety of receptor classes, including neurotrophin TrkA receptors^42^, where NGF binding to TrkA in the axonal terminals leads to receptor endocytosis. TrkA receptor on the signaling endosomes continues to be active and associates with downstream signaling components^42^. As another example, G-protein coupled receptors (GPCRs) can form signaling endosomes after binding to beta-arrestins^43–45^. Here, we observed that a significant portion of DMA-1 is internalized into endosomes in the PVD neuron. We have previously shown that DMA-1 interacts with TIAM-1, a Rac GEF that shows both diffuse and endosomal populations^46^. Thus, it is plausible that endosomal DMA-1 could directly recruit TIAM-1 to endosomes where it might recruit Rac to promote local actin polymerization^47^. However, we did not observe the robust presence of RAB-10 positive endosomes near or in outgrowing dendrites. Furthermore, a previous study showed that the GEF activity of TIAM-1 might not be required for its function in supporting PVD dendrite outgrowth and branching^22^.

Third, through this study, we uncovered a previously unknown function of endocytosis and recycling of membrane receptors. When DMA-1 is endocytosed, it dissociates from its bound transmembrane ligand SAX-7/L1CAM. Once it is reinserted on the plasma membrane through recycling endosomes, DMA-1 is now ligand-free and can diffuse into nascent filopodia, where it is ready for binding to available ligands. We have presented comprehensive experimental support for this model. We found three genetic manipulations (*kpc-1(null), hpo-30(R186A)* and *dma-1(C470Y)*) which reduce the endocytosis of DMA-1, causing DMA-1 to accumulate on the plasma membrane. In all three mutants, dendrite growth is severely blocked due to the absence of DMA-1 in the nascent filopodia. Furthermore, previous studies showed that the reinsertion of DMA-1 onto the membrane requires the small GTPase RAB-10 and the exocyst complex^31^. In *rab-10* mutants, DMA-1 and HPO-30 accumulate in intracellular vesicles and fail to be recycled onto the plasma membrane^30,31^, and *rab-10* mutants also exhibit severe PVD outgrowth defects^30,31^. Finally, when the endocytosis or recycling of DMA-1 is perturbed, DMA-1 becomes constitutively localized to the plasma membrane where it is immobilized through binding to its ligand (SAX-7/L1CAM). Together, these results suggest a population of diffusible DMA-1 on the membrane is required to promote dendrite outgrowth and branching. Because receptor endocytosis is a general phenomenon, it is plausible similar models might support branch morphogenesis of other dendrites and axonal terminals.

## Acknowledgements

We thank Liqun Luo for comments on the manuscript. Colleagues in the Shen laboratory for insightful discussions and comments. We thank the CGC for strains, which is funded by NIH Office of Research Infrastructure Programs (P40 OD010440), and the International C. elegans Gene Knockout Consortium. This work is supported by Howard Hughes Medical Institute and NIH NINDS 1R01NS082208 to K.S. and supported by the National Natural Science Foundation of China grants (31970919 and 31800861) to W.Z.

## Methods

### *C. elegans* strains and plasmids

(A) *C. elegans* animals were grown on nematode growth medium (NGM) plates using OP50 Escherichia coli as a food source and maintained according to standard procedure unless otherwise noted (Brenner, 1974). Before imaging, animals were raised at 20°C for at least one generation. N2 Bristol worms were used as the wild-type strain*. C. elegans* plasmids were generated in a pSMdelta vector backbone.

### EMS mutagenesis and forward genetic screens

For the *kpc-1(xr58)* genetic modifier screen, *kpc-1(xr58); wyIs592* worms were mutagenized at the L4 stage with 50mM ethyl methane sulfonate (EMS). F2 animals were screened under a fluorescent compound microscope for PVD morphology defects resembling *kpc-1* full loss-of-function mutants, including short and disorganized secondary dendrites. The *wy1008* allele was isolated from a screen from 20,000 haploid genomes and mapped to the *hpo-30* gene using standard SNP mapping^48^. The *wy1008* allele carries a missense C-to-T point mutation flanked by the sequences CAAAAGCT and TCTGGCCT, resulting in an E100K amino acid substitution.

To identify additional mutants defective in PVD dendrite morphogenesis, *wyIs594* (*ser2prom3::myr-gfp*) animals at the L4 stage were mutagenized with 50 mM EMS. Five F1 animals were transferred into individual plate seeded with OP50. About 100 F2 animals from each plate were examined under a fluorescent compound microscope for abnormal PVD dendrite arborization. The *zac98* allele was isolated from a screen which covered about 30,000 haploid genomes. *zac98* failed to complement the *dma-1* null allele *wy686*. The *zac98* allele carries a missense G-to-A point mutation flanked by the sequences AATGTCCGAT and CGCGACACCA, resulting in an C470Y amino acid substitution. The *zac227* allele contains a T to A (T1315A) mutation which affects the splicing of intron 5 of the *hpo-30* gene.

### CRISPR/Cas9 genome editing

Endogenous labeling of DMA-1 and RAB-10, as well as the generation of the *wy1220*, *wy1437*, and *wy1924* alleles were created by injection of CRISPR/Cas9 protein complexes into the gonad. Injection mixes consisted of 1.525 uM ALT-R Cas9 protein (IDT), 1.525 uM tracRNA (IDT), 1.525 uM crRNA (IDT), and repair templates consisting of 0.5 uM single-stranded DNA (IDT) or 0.15 −2 uM double-stranded PCR template. Single-stranded DNA repair templates were used for engineering the point mutations in the *wy1220* and *wy1437* alleles and were designed with a 30bp homology arm flanking the edited region. Double-stranded PCR repair templates were used for fluorophore insertions. All repair templates either mutated the protospacer adjacent motif (PAM) site or introduced silent mutations within the crRNA target sequence.

The *wy1437* allele was engineered using CRISPR/Cas9 to mimic a missense G-to-A point mutation resulting in a C470Y point mutation that was originally isolated by an EMS screen performed in Dr. Wei Zou’s lab. The *wy1924* allele was engineered using CRISPR/Cas9 to delete the LRR domain of DMA-1.

The following crRNAs were used:

*wy1616*: TGAAGAGCATGTCATACGGT

*wy1220*: TTTTACATAAATGGGTCCAA and GTTTTATCGAGAAGAGAACG

*wy1437*: GAATGACCTTCCACAAGCGATGG

*wy1924*: AGATCTAATACTTCCAAAGA and GTTGAACAAGTGTACGTAGA

### Sample preparation for Western blotting

For Western blots performed on S2 cell lysates: S2 cells were transfected with Pactin>hpo-30::gfp, Pactin>FLAG::hpo-30, or plasmids with R to A point mutations in Pactin>hpo-30::gfp (see Table S2). For cells grown in the presence of a furin inhibitor (Fig 4B), 50 uM Decanoyl-RVKR-CMK (Tocris) was added to the S2 media during transfection.

Three days after transfection, cells were lysed in RIPA buffer (Thermo Fisher) with 1x Halt Protease Inhibitor Cocktail (Thermofisher) for 30 minutes on ice. Cell lysates were spun at 13,000 rpm for 10 minutes at 4 degrees Celsius. Supernatants were collected for Western blot analysis in NuPAGE LDS Sample Buffer (Life Technologies) supplemented with DTT (GoldBio), and proteins were detected with a mouse antibody to GFP (1:2000, Roche), mouse antibody to FLAG (1:1000, Sigma), or mouse antibody to actin (1:5000, Abcam).

For Western blots performed on worm lysates: Worms were collected from one 10 cm NGM plate for each worm strain by washing with M9 buffer. Worms were spun at 2000 rpm and the worm pellet was washed with M9 1-3 times to remove residual bacteria. Worm pellets were lysed with an equal volume of NuPAGE LDS Sample Buffer (Life Technologies) supplemented with DTT (GoldBio), boiled for 10 minutes, then spun at 13,000 rpm for 10 minutes at room temperature. Supernatants were collected for Western blot analysis using a mouse antibody to GFP (1:2000, Roche) or mouse antibody to actin (1:5000, Abcam).

### Confocal imaging of *C. elegans*

All images were acquired at room temperature in live *C. elegans*. For images of mature PVD dendrite morphology, L4 and 1DOA stage worms were anesthetized using 10mM levamisole in M9 buffer and mounted on 3% agarose pads. Worms were then imaged on a 3i spinning disk microscope with a CSU-W1 spinning disk (Yokogawa) using a C-Apochromat 40x / 0.9 NA water immersion objective, or a spinning disk microscope with a CSU-X1 spinning disk (Yokogawa) using a 40x oil immersion objective. Images were acquired as z-stacks (0.75 um/step, 15-18 steps), and maximum-intensity projections were used for analysis and data presentation.

For images of endogenous DMA-1 or RAB-10 protein localization, worms were similarly anesthetized and mounted, then imaged on a spinning disk microscope with a CSU-X1 spinning disk (Yokogawa) using a 100x oil immersion objective. Single z-slices were chosen for all analyses.

For live time-lapse imaging of the developing PVD dendrite arbor and endogenous DMA-1:GFP or GFP::RAB-10 during PVD dendrite growth, L3 stage worms were mounted onto a glass-bottom imaging dish (MatTek)^49^. Worms were picked into a 5 mM droplet of levamisole and covered with a 5% agarose pad. Worms were then imaged on a spinning disk microscope with a CSU-X1 spinning disk (Yokogawa). For developing PVD dendrite arbor, images were acquired every 90 seconds as z-stacks (0.75 um/step, 12 steps) using a 40x oil objective and maximum-intensity z-projections were used for analysis and data presentation. For endogenous DMA-1::GFP during PVD dendrite outgrowth, images are captured utilizing a 100×/1.4 NA objective, 488 nm and 561 nm lasers, with a time interval of 2.5 seconds for a duration of up to 8 minutes. For endogenous GFP::RAB-10 during PVD dendrite outgrowth, images were acquired every 15 seconds as z-stacks (0.3 um/step, 6 steps) using a 63x oil objective and maximum-intensity z-projections were used for analysis and data presentation.

### Fluorescence recovery after photobleaching (FRAP) of DMA-1::GFP

FRAP assays were performed on a 3i spinning disk microscope with a CSU-W1 spinning disk (Yokogawa) using a C-Apochromat 63x / 1.2 NA water immersion objective with 2x magnification enhancer. Photobleaching was performed using a Vector x,y scanner (3i) and single z-slice images were collected every 15 seconds.

FRAP analysis was performed in ImageJ/Fiji. Images were cropped and registered using the StackReg plugin and corrected for photobleaching resulting from time-lapse imaging. FRAP intensity within the photobleached region was normalized to the average intensity of two background measurements taken before bleaching (100%), and the intensity immediately after bleaching (0%).

### Quantification of dendrite growth dynamics and enrichment of DMA-1::GFP in nascent filopodia

To measure the speed of dendrite growth, all the 2° dendrites that initiate protrusion from 1° branch were selected for quantification. Each growing secondary dendrite was tracked for 15 minutes after initiation. The frame captured just prior to the detection of the dendrite protruding from the 1° branch was designated as time 0 min.

To measure the enrichment of DMA-1::GFP in nascent filopodia, all the filopodia protrusions captured in the movies are selected for quantification. We define the frame right before the protrusion occurs as 0 sec. Lines were drawn along the nascent filopodia at timepoint 7.5 sec and their mother dendrite at 0 sec. Adjacent regions in the same frame were used as the background. Enrichment of DMA-1::GFP in nascent filopodia = (Mean intensity of nascent filopodia -Mean intensity of background) / (Mean intensity of mother dendrite -Mean intensity of background).

### Statistical analysis

All data are displayed as the mean ± standard error of the mean (SEM). Statistical comparisons were conducted using two-tailed Mann–Whitney tests (to test for differences between two groups) or Brown-Forsythe ANOVA with Dunnett’s multiple comparisons test (to test for differences between three or more groups). Sample sizes are indicated for each figure. ****, p < 0.0001; ***, p < 0.001; **, p < 0.01; *, p < 0.05 in all graphs. All statistical analyses and graph construction were performed using Prism 9.0 software (GraphPad Software, Inc)

---

**Figure S1. KPC-1 genetically interacts with HPO-30**

(A) Representative images of PVD dendrite morphology in wild-type, *kpc-1(null),* and *kpc-1(xr58)* partial loss-of-function animals at the L4 stage. (B) Representative images of PVD dendrite morphology in *kpc-1(xr58); hpo-30(wy1008)* double mutants, *hpo-30(wy1008)* single mutants, and *hpo-30(null)* animals at the L4 stage. (C) Quantification of number of quaternary branches in a region 150 µm anterior to the PVD cell body, for genotypes shown in A and B. (D) Quantification of percentage of extended secondaries for genotypes shown in A and B. Extended secondaries were measured as the percentage of secondary branches per worm that extended at least halfway to the tertiary dendrite line. For (C and D), n values within each bar, error bars indicate the standard error of proportion. Statistical comparison was performed using Brown-Forsythe ANOVA with Dunnett’s multiple comparisons test.

**Figure S2. Uncleavable HPO-30(R184A) binds to DMA-1 and HPO-30 mislocalises to the plasma membrane in *kpc-1(null)* mutants**

(A) Western blot against GFP in cell lysates from Drosophila S2 cells expressing wild-type HPO-30::GFP or HPO-30::GFP with individual arginine residues mutated to alanine as indicated. Top schematic represents the location of the R186A mutation in the second extracellular loop of HPO-30. (B) Western blots showing co-IP between DMA-1::HA and HPO-30(WT)::GFP or HPO-30(R186A)::GFP expressed in S2 cells. Samples were immunoprecipitated with an anti-GFP nanobody. (C) Representative images of HPO-30::GFP in wild-type and *kpc-1(null)* animals. Yellow rectangles indicate locations of magnified views on the right.

**Figure S3. Endosomal DMA-1 is required for PVD dendrite morphogenesis**

(A) Representative images of PVD morphology in wild-type and *dma-1(C470Y)* mutants at the L4 stage. (B) Quantification of number of quaternary branches in a region 150 µm anterior to the PVD cell body for *dma-1(C470Y)* animals. (C) Quantification of percentage of extended secondaries in *dma-1(C470Y)* mutants. In (B and C), n values within each bar, error bars indicate the standard error of proportion. Statistical comparison was performed using Brown-Forsythe ANOVA with Dunnett’s multiple comparisons test. (D) Endogenous DMA-1::GFP knock-in in *dma-1(C470Y)* animals. Yellow rectangles indicate locations of magnified views on the right. Arrowheads indicate punctate structures representing putative endosomes. (E) Colabeling of endogenous DMA-1::GFP with endogenous mScarlet::RAB-10 (mScarlet::RAB-10; expressed cell-specifically in PVD) in the cell bodies of *dma-1(C470Y)* mutant animals. (F) Colabeling of endogenous DMA-1::GFP with endogenous mScarlet::RAB-10 (mScarlet::RAB-10; expressed cell-specifically in PVD primary dendrite of wild-type and *dma-1(C470Y)* animals. Yellow arrowheads indicate locations of mScarlet::RAB-10 puncta. (G and H) Quantification of the percentage of mScarlet::RAB-10 puncta that overlap with DMA-1::GFP puncta in the cell bodies (G) and primary dendrites (H) in *dma-1(C470Y)* animals. (I) Quantification of DMA-1::GFP fluorescence intensity measured outside of mScarlet::RAB-10 puncta in *dma-1(C470Y)* animals. For (G, H and I), n values within each bar, error bars indicate the standard error of proportion. Statistical comparison was performed using Brown-Forsythe ANOVA with Dunnett’s multiple comparisons test.

**Figure S4. Removal of SAX-7 in *kpc-1(null), hpo-30(R186A)*, or *dma-1(C470Y)* animals increases dendritic growth**

(A) Representative images of PVD dendrite morphology in indicated genotypes at the L4 stage. (B) Quantification of the mean length of all secondary dendrites in individual worms. n values within each bar, error bars indicate the standard error of proportion. Statistical comparison was performed using Brown-Forsythe ANOVA with Dunnett’s multiple comparisons test.

**Figure S5. Delayed enrichment of DMA-1::GFP in the nascent filopodia of *hpo-30(R186A)* and *dma-1(C470Y*) mutants**

Time-lapse imaging of developing PVD dendrites with DMA-1::GFP (green) and PVD marker (*ser2prom3::mCherry* (magenta)) in *hpo-30(R186A)*, *dma-1(C470Y)* or *sax-7(null)* animals. Arrowheads point to the tip of outgrowing dendritic branches.

**Figure S6. Endogenous GFP::RAB-10 is not localized in the growing PVD dendrites**

Time-lapse imaging of developing PVD dendrites with GFP::RAB-10 (green) and (*ser2prom3::mCherry* (magenta)) in wild-type animals. Blue arrows indicate growing dendrites. Yellow arrowheads indicate GFP::RAB-10 puncta.

**Figure S7. Stereotyped dendritic arbors requires the recycling of ligand-free DMA-1 through HPO-30 cleavage and selective stabilization through ligand-bound DMA-1**

Proper dendritic morphogenesis requires rounds of dendrite outgrowth and stabilization. Dendrite stabilization requires the binding of ligands to the HPO-30/DMA-1 complex. Dendritic outgrowth is dependent on endocytic recycling of DMA-1 to generate mobile ligand-free DMA-1. The activation of DMA-1 recycling requires ligand-binding of DMA-1 to its ligand and cleavage of HPO-30 by KPC-1. MNR-1 is drawn in dark green. SAX-7 is drawn in purple. LECT-2 is drawn in red. DMA-1 is drawn in brown. HPO-30 is drawn in yellow.

**Table S1.** Worm strains used in this study.

| Strain Name | Genotype | Description |
| --- | --- | --- |
| TV15911 | *wyls592 III* | *ser2prom3::myr-gfp; Podr-1::rfp* |
| TV17248 | *sax-7(nj48) IV; wyls592 III<sup>3</sup>* | Referred to as *sax-7(null)* |
| TV29097 | *dma-1(wy1924) I; wySi919 V* | *wy1924 = dma-1(ΔLRR)*, *wySi919 = Pdes-2::myr-mScarlet::let-858* |
| TV24863 | *hpo-30(zac227) V; wyls592 III* | *zac227 = hpo-30(ΔICD)* |
| TV29380 | *hpo-30(zac227) V; dma-1(wy1924) I; wySi919 V* |  |
| TV29319 | *hpo-30(zac227) V; dma-1(wy908) I; wyls592 III* | *wy908 = dma-1(ΔICD)* |
| TV17465 | *dma-1 (wy908) I; wyls592 III* |  |
| TV29856 | *kpc-1(gk8) I; wyls592 III* | Referred to as *kpc-1(null)* |
| TV17268 | *kpc-1(xr58) I; wyls592 IIII* |  |
| TV19400 | *kpc-1(xr58) I; hpo-30(wy1008) V; wyls592 III* |  |
| TV19079 | *hpo-30(wy1008) V; wyls592 III* |  |
| TV16821 | *hpo-30(ok2047) V; wyls592 III* | Referred to as *hpo-30(null)* |
| TV24410 | *hpo-30(wy1220) V; wyls592 III* | *wy1220 = hpo-30(R186A)* |
| TV24913 | *dma-1(wy1246) I* | *wy1246 = dma-1::GFP* (Endogenously tagged) |
| TV27036 | *dma-1(wy1246) I; kpc-1(gk8) I; wyls581* | *wyls581 = ser2prom3::myr-mCherry + Podr-1::gfp* |
| TV27089 | *dma-1(wy1246) I; hpo-30(wy1220) V* |  |
| TV27759 | *dma-1(wy1246) I; rab-10(wy1616) I; wyls910 X* | *wyls910 = ser2prom3::FLP + Punc-122::bfp*<br>*wy1616 = flp-on mScarlet::RAB-10* |
| TV27781 | *dma-1(wy1246) I; rab-10(wy1616) I; kpc-1(gk8) I; wyls910 X* |  |
| TV27863 | *dma-1(wy1246) I; rab-10(wy1616) I; hpo-30(wy1220) V; wyls910 X* |  |
| TV27037 | *dma-1(wy1246) I; sax-7(nj48) IV* |  |
| TV26091 | *dma-1(wy1437) I* | *wy1437 = dma-1(C470Y)::GFP*<br>(Endogenously tagged) |
| TV27649 | *dma-1(wy1246) I; rab-10(wy1616) I; sax-7(nj48) IV; wyls910 X* |  |
| TV27865 | *dma-1(wy1437) I; rab-10(wy1616) I; wyls910 X* |  |
| TV26828 | *dma-1(zac98) I; wyls592 III* | *zac98 = DMA-1(C470Y)* |
| TV17247 | *dma-1(tm5159) I; wyls592 III* | Referred to as *dma-1(null)* |
| TV25655 | *hpo-30(wy1220) V; wyls738; wyls592 III* | *wyls738 = ser2prom3::dma-1:gfp + Podr-1::gfp* |
| TV21180 | *hpo-30(ok2047) V; wyls738; wyls592 III* |  |
| TV23577 | *wyEx9544; hpo-30(ok2047) V; wyls592 III* | *wyEx9544 = ser2prom3::hpo-30(R186A)::gfp; Pmyo-2::mCherry (Line 1)* |
| TV23578 | *wyEx9545; hpo-30(ok2047) V; wyls592 III* | *wyEx9545 = ser2prom3::hpo-30(R186A)::gfp; Pmyo-2::mCherry (Line 2)* |
| TV25343 | *wyEx10069; hpo-30(ok2047) V; wyls587* | *wyEx10069 = ser2prom3::hpo-30:gfp; Pmyo-2::mCherry* |
| TV25344 | *wyEx10070; hpo-30(ok2047) V; wyls587* | *wyEx10070 = ser2prom3::hpo-30(R186A)::gfp; Pmyo-2::mCherry* |
| TV27592 | *dma-1(wy1437) I; hpo-30(ok2047) V* |  |
| TV27373 | *dma-1(wy1437) I; sax-7(nj48) IV* |  |
| TV18815 | *wyEx7786; qyls369* | *wyEx7786 = ser2prom3::hpo-30cDNA::mCherry; Podr-1::gfp*<br>*qyls369 = ser2prom3::dma-1::gfp; unc-119 (+)* |
| TV27864 | *dma-1(wy1246) I; rab-10(wy1616) I; hpo-30(ok2047) V; wyls910 X* |  |
| TV27065 | *rab-10(wy1298) I; wyls581 IV; wyls910 X<sup>29</sup>* | *wy1298 = FRT::GFP::FRT::rab-10*<br>This strain uses the FLP-on system to cell-specifically tag RAB-10 in the PVD. |

**Table S2.**
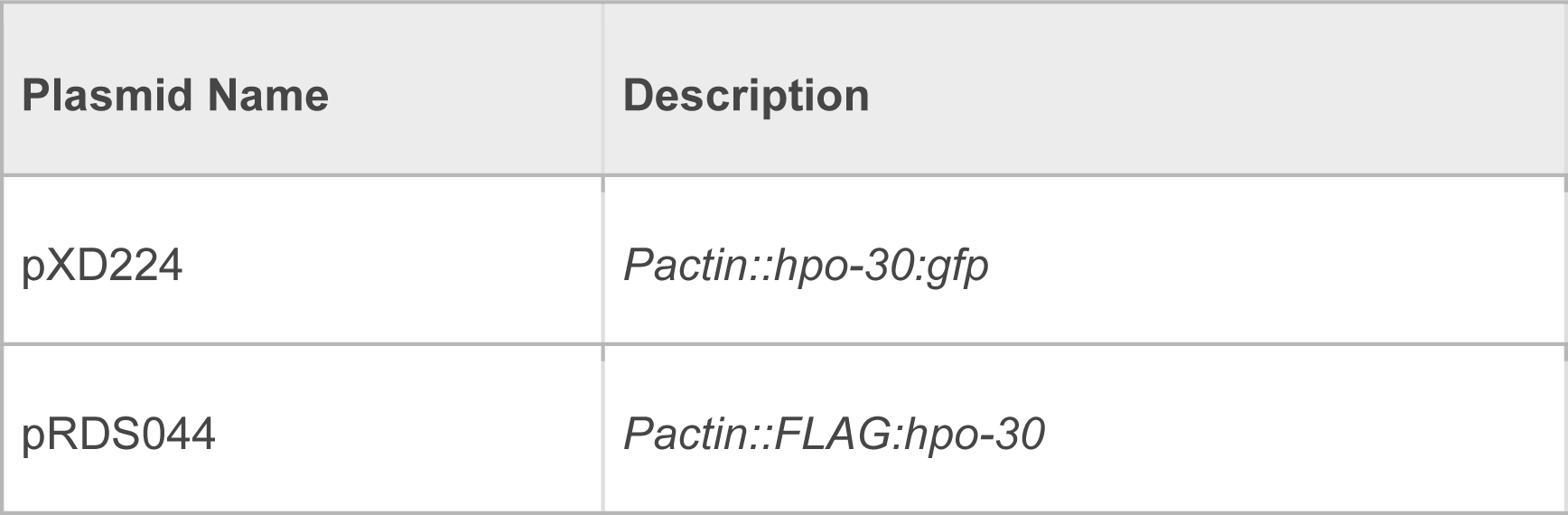
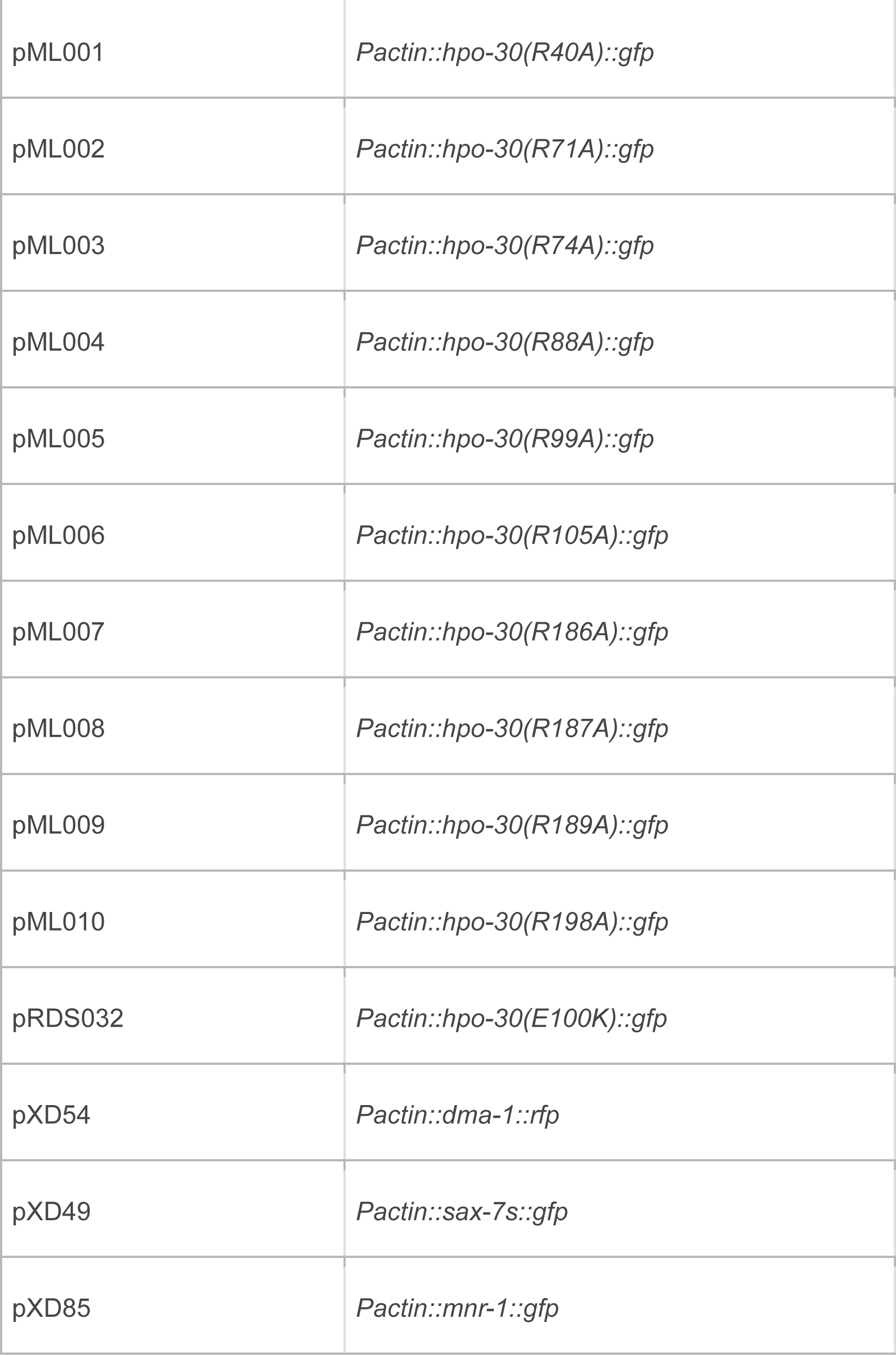
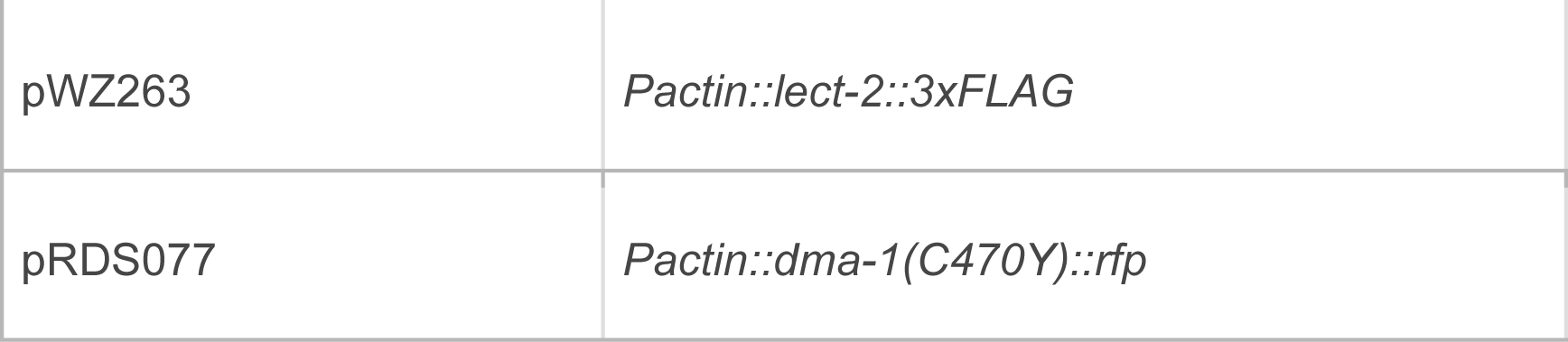
S2 cell plasmids used in this study.

